# Experimental depletion of gut microbiota diversity reduces host thermal tolerance and fitness under heat stress in a vertebrate ectotherm

**DOI:** 10.1101/2021.06.04.447101

**Authors:** Samantha S. Fontaine, Patrick M. Mineo, Kevin D. Kohl

## Abstract

Predicting the responses of ectotherms to climate change is a global conservation priority which requires identifying factors that influence how animals respond physiologically to changing temperature. Host-associated microbial communities impact animal physiology and have been shown to influence host thermal tolerance in invertebrate systems. However, the role of commensal microbiota in thermal tolerance of ectothermic vertebrates is unknown. Here we show that experimentally depleting the diversity of the tadpole gut microbiome through environmental water sterilization reduces the host’s acute thermal tolerance to both heat and cold, alters the thermal sensitivity of locomotor performance, and reduces animal survival under acute heat stress. We show that these tadpoles have reduced activities of mitochondrial enzymes and altered metabolic rates compared to tadpoles colonized with a diverse microbiota, which could underlie differences in thermal phenotypes. Our results demonstrate, for the first time, a link between the gut microbiome of an ectothermic vertebrate and the host’s thermal tolerance, performance, and fitness, thus highlighting the importance of considering host-associated microbial communities when predicting species’ responses to climate change.

## Main

Temperature is a crucial environmental factor impacting the physiological performance and fitness of ectothermic animals, which, are expected to be especially vulnerable to the deleterious effects of global climate change^1, 2^. Indeed, global surveys have shown declines in several ectothermic vertebrate groups due to impacts of climate change^3, 4^. Continuing to predict how these animals will physiologically respond to changing temperature regimes is a global conservation priority which requires a thorough understanding of the mechanisms that contribute to host thermal tolerance and the impact of temperature on whole-animal performance^5^. Decades of research have assessed the thermal tolerance of ectotherms under a range of conditions by quantifying metrics such as the critical thermal minimum and maximum (CT_min_ and CT_max_), performance across temperatures (thermal performance curves), and survival at sublethal temperatures^6–8^. Such studies have revealed that while thermal tolerance is in part a genetic trait^9^, it is also plastic, and can be shaped by environmental factors, such as time of day^10^, season^11^, oxygen availability^10^, feeding^12^, and infection status^13, 14^.

Host-associated microbial communities, or microbiomes, have recently emerged as a critical factor that regulate how animals respond to their external environments through impacts on host physiology (e.g. digestion, immune function, energy metabolism) and gene expression^15–17^, and may therefore also play a role in host thermal tolerance. However, we currently lack a complete understanding of this relationship across ectotherm groups. In invertebrate animal systems, there is indeed a relationship between symbiotic microbes and host thermal tolerance. For example, in sea anemones, corals, and aphids, acquisition of particular symbionts can buffer against the lethal effects of heat shock, allowing hosts to persist in otherwise extreme habitats^18–21^. Further, axenic flies exhibit decreased survival under heat stress compared to conventional flies^22^, and microbial transplants can alter fly thermal tolerance^23^.

While there have been no manipulative studies, there is correlative evidence that host-associated microbial communities may also impact thermal tolerance in ectothermic vertebrate species. Environmental temperature influences the diversity and composition of the gut microbiome in several systems^24–28^. In lizards, exposure to heat alters the gut microbiome, some aspects of microbial composition are associated with the critical thermal maximum of hosts^29^, and loss of microbial diversity is correlated with reduced animal survival^27^. Additionally, artificial selection for cold tolerance in fish alters microbiome composition and sensitivity to cold exposure, suggesting a relationship between these phenotypes^30^. However, a major challenge of the microbiome field is to move past correlations between the microbiome and host phenotypes, and to truly demonstrate functional effects of the microbiome on host biology^31^.

Here, we use a series of manipulative experiments to test the hypothesis that the host-associated microbiome influences thermal tolerance in an ectothermic vertebrate. We demonstrate, for the first time, a direct relationship between the gut microbiome of a vertebrate ectotherm and the host’s thermal tolerance, performance, and fitness. Specifically, we find that tadpoles with an experimentally depleted gut microbial community exhibit a significant reduction in tolerance under acute thermal stressors, impaired fitness under prolonged heat stress, and a change in the thermal sensitivity of whole-organism performance compared to those with a more diverse microbial community. To enhance our understanding of the physiological integration of host-microbe interactions across levels of biological organization^32, 33^, we compare several physiological metrics across microbiome treatment groups, including cell membrane phospholipid composition, mitochondrial enzyme activities, and metabolic rate, which are traits associated with thermal tolerance in ectotherms^6, 34, 35^. Collectively, our study demonstrates that host associated-microbial communities are an important factor contributing to the thermal physiology of ectothermic vertebrates on several scales, from the subcellular to whole-organism level, which is important information given the global declines in ectotherm diversity and the continued threat of global climate change.

## Results and Discussion

### Experimental depletion of environmental microbiota reduced tadpole gut microbiome diversity and altered composition

For all experiments, we used laboratory-reared tadpoles from wild adults of the green frog *(Lithobates clamitans).* Green frogs are an abundant and widespread anuran amphibian across eastern North America, and can adjust their physiology in response to temperature during both larval and adult lifestages^36, 37^. To manipulate the gut bacterial community, tadpoles were divided into two treatment groups: colonized and depleted. Colonized tadpoles were raised in autoclaved laboratory water seeded with 25% natural pond water from their parent’s site of capture to provide microorganisms for colonization, whereas depleted tadpoles were raised in autoclaved laboratory water seeded with 25% autoclaved pond water, to reduce microorganisms available for colonization^38^.

In our first experiment, we raised colonized and depleted tadpoles spawned from adults collected from Louisiana (USA) at three acclimation temperatures: 14°C, 22°C, and 28°C, and used high-throughput 16S rRNA amplicon sequencing to compare gut bacterial communities across treatment groups. Additionally, we tracked the bacterial communities of our stored pond water (kept at 4°C) weekly during the experiment and compared these communities to samples collected fresh from the pond to determine how well our colonized treatment reflected natural pond communities. Water samples (fresh and stored) had distinct bacterial community composition from tadpole gut samples (Supplementary Figure 1a; PERMANOVA, pseudo-F= 8.30, p<0.001). Additionally, stored water samples had a significantly different bacterial community composition than fresh water samples (Supplementary Figure 1b; PERMANOVA, pseudo-F= 1.91, p=0.03). While fresh and stored water samples may have differed based on amplicon sequence variant (ASV)-level comparisons, at a phylum level, the community composition was largely the same and was dominated by Proteobacteria, Acidobacteria, Bacteroidetes, and Actinobacteria (Supplementary Figure 1c). Our results are consistent with other studies finding relative stability in microbial communities of pond water stored at 4°C in the laboratory^39^. In tadpole gut bacterial communities, the dominant phyla overall were Firmicutes and Proteobacteria (Supplementary Figure 2), which appear to be common commensals of larval amphibians, having been documented in high abundance in the guts of several other tadpole species^40, 41^, as well as in the early life-stages of the eastern newt^42^.

Acclimation temperature significantly altered the tadpole gut microbiome such that tadpoles raised at warmer temperatures had less diverse bacterial communities than those raised at cooler temperatures (Fig. 1a, Supplementary Figure 3a, Supplementary Table 1; GLMM, p<0.001), and bacterial community composition and variability were distinct across all temperature groups (Fig. 1b, Supplementary Figure 3e-f, Supplementary Table 1; PERMANOVA, p<0.001; PERMDISP, p<0.01). A single bacterial phylum, Acidobacteria, had a relative abundance that was associated with acclimation temperature, and was most abundant in the 14°C group (Supplementary Table 2; MaAsLin, corrected p=0.03). Our results are consistent with those that find reduced gut bacterial diversity and altered composition at increased temperatures in various ectotherms^23, 24, 27, 28, 43, 44^. The causes of these changes are unknown but may involve temperature-dependent growth of bacterial communities, or indirect effects of temperature on host physiology and/or behavior^24, 27^.

**Fig. 1.**
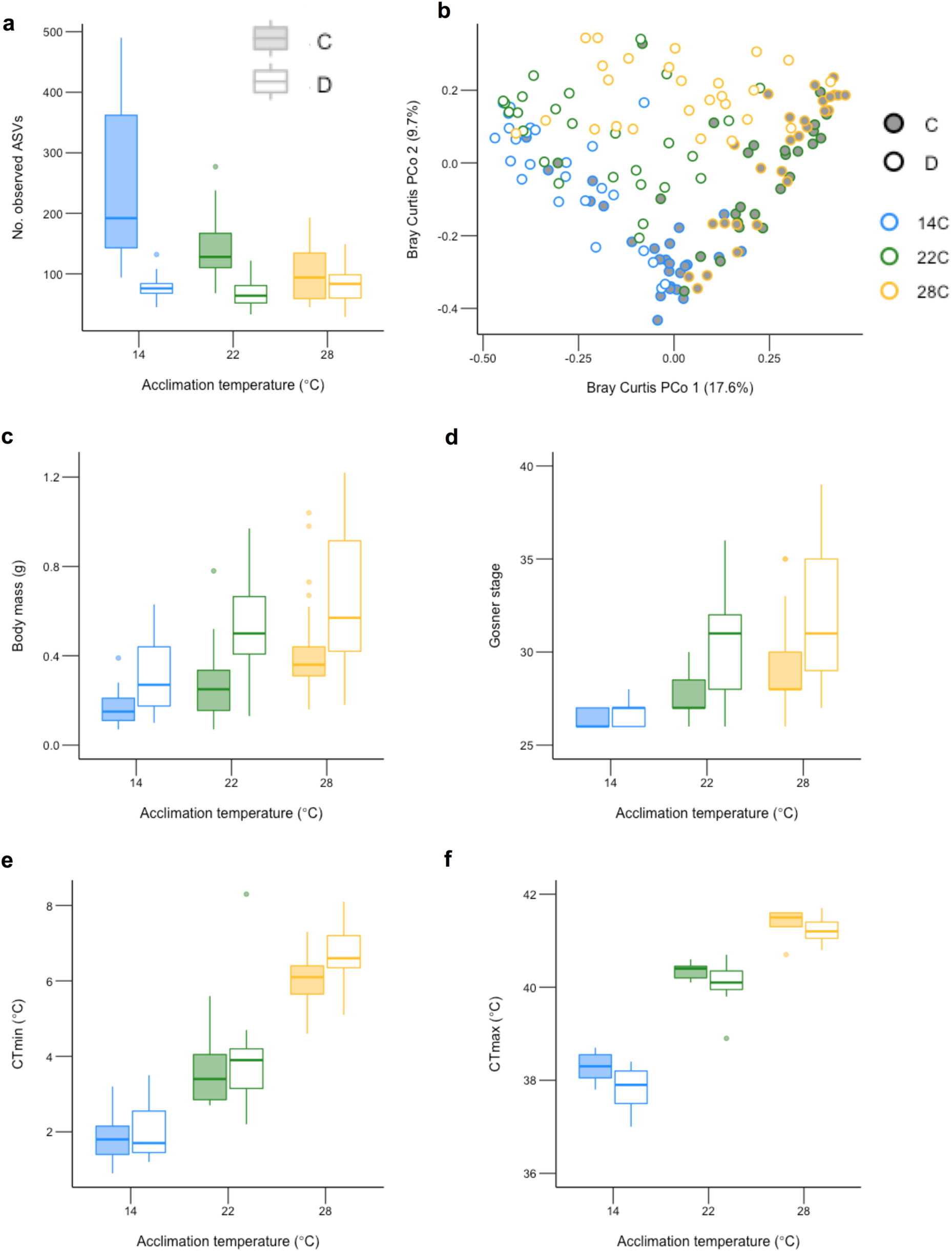
Effects of microbial colonization treatment and acclimation temperature on tadpole gut microbial communities, morphometrics, and acute thermal tolerance. **a,** Number of observed bacterial amplicon sequence variants (ASVs) in tadpole gut microbial communities **b,** Principal Coordinate (PCo) analysis plot based on Bray-Curtis dissimilarity of gut microbial communities between samples. Percentages represent the proportion of variation explained by each axis **c,** tadpole body mass **d,** tadpole developmental stage based on the Gosner system, **e,** tadpole critical thermal minimum (CT_min_) f, tadpole critical thermal maximum (CT_max_). For boxplots **a,c,** and **d,** N=27 animals per group. For boxplots **e and f**, N=11 animals per group. In each boxplot, the center line represents the median, the length of the box extends through the IQR, and whiskers extend to 1.5x IQR. All points outside this range are plotted individually. C= colonized tadpoles and D= depleted tadpoles. Colors represent tadpole acclimation temperature.

We found that depleted tadpoles had reduced gut bacterial diversity (Fig. 1a, Supplementary Figure 3a-c, Supplementary Table 1; GLMM, p<0.001), and this effect interacted with temperature (Fig. 1a, Supplementary Figure 3a-c, Supplementary Table 1; GLMM, p<0.05). Because increasing temperatures reduced bacterial diversity even in colonized individuals, tadpoles at the coolest acclimation temperature exhibited larger differences in microbial diversity between colonized and depleted animals than those at warmer acclimation temperatures. Depleted tadpoles also harbored bacterial communities with a distinct composition compared to colonized tadpoles (Fig. 1b, Supplementary Figure 3e-f, Supplementary Table 1; PERMANOVA, p<0.001). Our results mirror those of a previous study that found raising tadpoles in sterilized water reduces the diversity and alters the composition of the gut bacterial community^38^. Additionally, depleted tadpoles exhibited greater degrees of intraindividual variability within their gut microbial communities than colonized tadpoles (Supplementary Table 1; PERMDISP, p<0.01). This difference could be due to development in a more stressful environment, as reduced diversity of microbial communities has been associated with increases in host stress hormones^45^, which can then reduce a host’s ability to regulate their commensal microbiome, and ultimately lead to greater stochasticity in the community^46^. Additionally, development in an environment with fewer microbial members may also increase community stochasticity due to ecological drift^47^. Within these communities, the relative abundances of seven bacterial phyla and 11 bacterial genera were impacted by microbial colonization treatment (Supplementary Table 2; MaAsLin, corrected p<0.05). The affected phyla included those such as Dependentiae, Plantomycetes, WPS-2, and Verrucomicrobia, which are common to aquatic and soil habitats^48–51^, and were also highly abundant in tadpole guts colonized with natural pond water, but were absent or present at very low abundances in depleted tadpoles (Supplementary Table 2). Because green frog tadpoles develop externally to the mother and lack parental care, it is likely a high proportion of their symbiotic microbes come from transmission of environmental microbes^52^, and therefore, depleting these communities results in a depleted gut microbiome.

However, neither microbial colonization treatment nor acclimation temperature impacted the total abundance of bacterial cells in the gut based on flow cytometry analyses (Supplementary Figure 3d; LME, p>0.05). This result supports the idea that there may be host species-specific carrying capacities that set limits to gut microbial population densities, regardless of differences in community composition^53^. Further, this result suggests that opposed to germ-free studies that compare colonized animals with those under the highly artificial state of sterility, our results may more accurately represent dynamics that occur in animal microbiomes in the wild. Specifically, in wild populations, microbes are consistently available for colonization, but anthropogenic or environmental impacts may alter their community diversity or composition^54^.

### Experimental depletion of microbiota altered tadpole development, morphometrics, and acute thermal tolerance

In our first experiment, after tadpoles had developed for seven weeks post-hatching, their acute thermal tolerance was measured via the critical thermal minimum (CT_min_) and maximum (CT_max_). We used Hutchison’s dynamic method^55^, in which temperatures are increased or decreased until a loss of the righting response occurs, to measure CT_min_ and CT_max_ in 11 individuals from each of the six treatment groups (acclimation temperature x microbial colonization). All remaining individuals (N = 16 animals per group) were euthanized for morphometric data without measuring thermal tolerance. In each tadpole, we measured body mass (g), body length (mm), body width (mm), Gosner stage^56^, and facial symmetry. To measure facial symmetry, the absolute value of the difference between the distance between each eye and the tip of the nose was measured. Greater degrees of asymmetry are associated with reduced developmental stability and stressful conditions^57^.

We found that temperature impacted tadpole growth and development such that tadpoles raised at warmer acclimation temperatures were significantly larger and more developed than tadpoles raised at cooler temperatures (Fig. 1c-d; Supplementary Figure 4a-b; Supplementary Table 3; GLMM, p<0.001). This finding was expected based on the wealth of studies demonstrating the positive relationship between environmental temperature and tadpole growth and development at sublethal temperatures^58–61^. We also observed that tadpoles raised at warmer temperatures exhibited decreased facial symmetry compared to those at cooler temperatures (Supplementary Figure 4c; Supplementary Table 3; GLMM, p<0.001) indicating our higher developmental temperatures may have increased tadpole stress and developmental instability.

We also found that depleted tadpoles were significantly larger and more developed than colonized tadpoles (Fig. 1c-d; Supplementary Figure 4a-b; Supplementary Table 3; GLMM, p<0.001). Specifically, depleted tadpoles were 55% larger in terms of body mass, and more developed by an average of two Gosner stages than colonized tadpoles, regardless of acclimation temperature. Our results are similar to studies in wild, laboratory, and agricultural systems that find enhanced growth and body size in animals treated with antibiotics to deplete gut bacterial communities^62–65^. There are several hypotheses that may explain this phenomenon. For example, animals invest heavily in their immune systems to both defend against pathogens, and regulate populations of commensal bacteria^66^, and thus, reducing gut microbial diversity may allow animals to invest more in growth and development as opposed to immune system regulation^65^. Interestingly, the gut microbiome of colonized tadpoles had higher proportions of the bacterial phylum Chlamydiae than depleted tadpoles (Supplementary Table 2), which are obligate intracellular pathogens known to exploit host nutrients and elicit an immune response^67^. Additionally, stress hormones are known to stimulate growth and development in tadpoles^68^, and reductions in microbial diversity can increase animal stress^45^. However, we did not measure physiological markers associated with stress, such as glucocorticoids, in our system, and we did not observe a significant effect of microbial colonization treatment on tadpole facial symmetry (Supplementary Figure 4c; Supplementary Table 3; GLMM, p>0.05).

We found that acclimation temperature predictably impacted tadpole thermal tolerance, such that tadpoles raised at warmer temperatures were more tolerant to heat, and tadpoles raised at cooler temperatures were more tolerant to cold. Specifically, animals raised at the coolest temperature (14°C) had a CT_min_ that was on average 4.5°C lower than animals raised at the warmest temperature (28°C) (Fig. 1e; GLMM, χ^2^= 182.04, p<0.001). Animals raised at the warmest temperature had a CT_max_ that was on average 3.3°C higher than animals raised at the coolest temperature (Fig. 1f; GLMM, χ^2^= 673.25, p<0.001). These results are consistent with studies demonstrating that many anuran amphibian species, including the green frog, can adjust their critical thermal limits adaptatively in response to acclimation temperatures^37^.

Notably, we observed that depleted tadpoles exhibited reductions in their tolerance to cool temperatures compared to colonized tadpoles. Specifically, these animals had higher CT_min_ values than colonized tadpoles after controlling for the effects of acclimation temperature, body size, and Gosner stage (Fig. 1e; GLMM, χ^2^= 5.30, p=0.02). Other studies demonstrating a relationship between gut microbiota and host cold tolerance are scarce in ectothermic systems, however, selection for cold tolerance in fish does alter the gut microbial community^30^. In mammalian systems, changes occurring in the gut microbiota upon exposure to cold ultimately lead an increased host tolerance to cold through enhanced energy acquisition^69, 70^. The mechanisms underlying tadpole tolerance to cold likely differ, as studies in mice demonstrate that microbial products facilitate host maintenance of internal body temperature during cold exposure, which would not be important in ectothermic systems. Regardless, our result warrants future study and may impact animals in the wild, as green frogs often overwinter as tadpoles and can be exposed to near freezing temperatures^36^.

Depleted tadpoles were also less tolerant to heat than their colonized counterparts. At each acclimation temperature, depleted tadpoles had lower CT_max_ values than colonized tadpoles (Fig. 1f; GLMM, χ^2^= 18.60, p<0.001). These results echo those of studies focused on invertebrate ectotherms that have shown microbial symbionts can enhance host heat tolerance^18–20, 23, 71^. The reduction in heat tolerance in microbially depleted tadpoles is not simply a function of the difference in growth and development between the two groups. Body mass and Gosner stage were included as covariates in the above models and did not change the significant relationship between microbial colonization and host thermal tolerance. Further, we observed a significant positive relationship between animal body mass and Gosner stage, and CT_min_ and CT_max_ (GLMM, χ^2^= 4.23 and 13.90 for body mass and Gosner stage respectively with CT_min_, and χ^2^= 4.06 and 17.88 for body mass and Gosner stage respectively with CT_max_, p<0.05 for all). For heat tolerance, this trend is the opposite of what would be expected if these factors were driving the relationship between thermal tolerance and microbial colonization treatment.

Taken together, our results show that depleted tadpoles have both a higher CT_min_ and lower CT_max_ than colonized tadpoles, resulting in a smaller overall window of thermal tolerance in these animals. Our study is the first to report a direct link between host associated-microbial communities and host thermal tolerance in an ectothermic vertebrate. Wider thermal tolerance breadths are often observed in populations that experience greater degrees of thermal variability^72^, and can be important in adapting to fluctuating environments^73^. Our results suggest that loss of gut microbial community diversity may hinder the ability of organisms to cope with temperature fluctuations in their environment that they may experience due to changes in global climate. Host-associated bacterial diversity may be lost due to many anthropogenic environmental impacts^54^, however, increasing temperatures themselves can also reduce gut microbial community diversity of ectotherms as evidenced by our results and others. Thus, negative impacts of climate warming may compound such that warming-induced losses of bacterial diversity may leave animals even more vulnerable to further warming due to potential reductions in their thermal tolerance.

### Experimental depletion of microbiota reduced tadpole survival under heat stress

We conducted an additional study addressing the differences in heat tolerance observed between groups, given that climate warming is a major threat to species worldwide^74^, as well as in our study area^75^. Therefore, in a second experiment, we sought to determine if depleted tadpoles, which experienced a significant reduction in their acute heat tolerance, would also incur fitness costs upon exposure to sublethal heat stress for longer time periods. Using adult frogs and pond water from a different geographic location (PA, USA), we raised colonized and depleted tadpoles at a single acclimation temperature of 22°C for seven weeks. In a subset of these tadpoles, we verified our results from experiment 1 to ensure consistency across populations. Using 16S rRNA amplicon sequencing, we again demonstrated that compared to colonized tadpoles, the gut microbiota of depleted tadpoles was different in community composition (Supplementary Figure 5d; Supplementary Table 1; PERMANOVA, p<0.001), exhibited a greater degree of intraindividual variability (Supplementary Table 1; PERMDISP, p<0.05) and was less diverse (Supplementary Figure 5a-b; Supplementary Table 1; Kruskal-Wallis, p<0.001), although in this set of animals, depleted tadpoles exhibited greater gut microbial community evenness than colonized tadpoles (Supplementary Figure 5c; Supplementary Table 1; Kruskal-Wallis, p<0.01). Additionally, the depleted tadpoles were again larger and more developed (Supplementary Figure 6a-c; Supplementary Table 3; GLM, p<0.05), and less heat tolerant (in terms of CT_max_) than colonized tadpoles (Supplementary Figure 6d; GLM, t= 2.23, p=0.03). The repeatability of our results demonstrates the impacts of microbial community colonization on these phenotypes are consistent regardless of population of origin.

After this verification, half of the remaining tadpoles in each microbial colonization treatment group were exposed to an increased temperature of 32°C for ten days, which was then increased to 34°C for another ten days, and finally increased to 36°C for a final ten days. The second half of the tadpoles remained at 22°C. We monitored survival of each individual daily for the full 30 days. We found that both temperature and microbial colonization strongly influenced tadpole survival. There was 100% survival in both microbial treatment groups at 22°C, and more variable survival across temperatures in the heat stress treatments (Fig. 2; discrete-time logistic regression, z= 4.79, p<0.001). However, colonized animals exhibited significantly greater survival than depleted animals under heat stress conditions (Fig. 2; discrete-time logistic regression, z= 2.07, p=0.04). Based on hazard functions from the logistic regression model, the risk of mortality in the depleted group was ∼5 times greater than in the colonized group at both 32 and 34°C, while there was 100% mortality in both groups at 36°C.

**Fig. 2.**
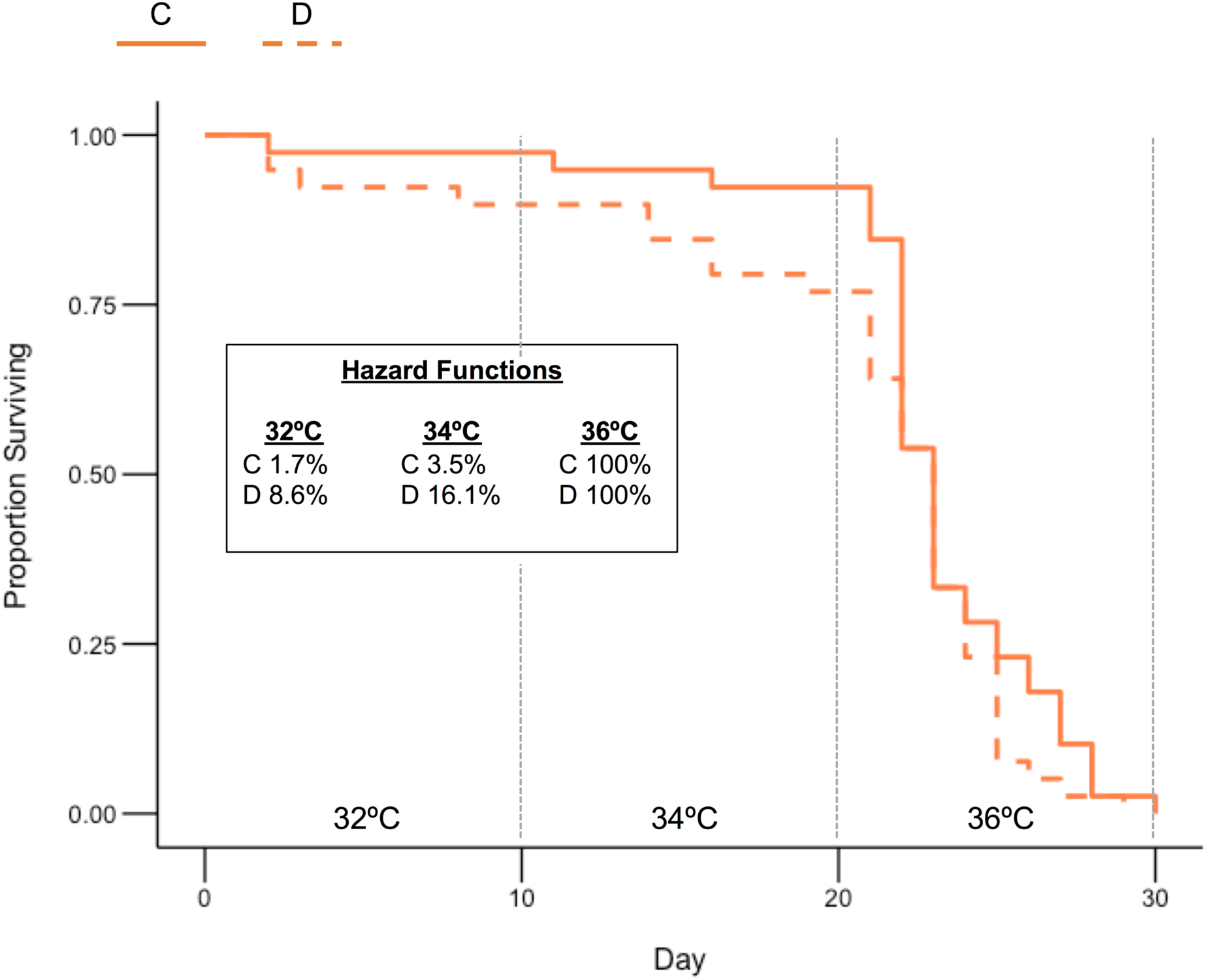
Survival of colonized and depleted tadpoles under heat stress conditions. Labels above the x-axis correspond to the temperature treatment at each timepoint of the experiment. Hazard functions were calculated from a discrete-time logistic regression and represent the risk of mortality of colonized and depleted tadpoles at each temperature. C= colonized tadpoles and D=depleted tadpoles. N = 39 animals per group at the beginning of the trial. A 22°C control group was not included in the analysis because there was 100% survival in this group.

Our results add to a growing body of literature demonstrating the importance of the host-associated microbiome in maintaining host thermal tolerance. In *Drosophila*, axenic flies exhibit poorer survival compared to conventional flies at lower heat stress temperatures (35-36°C), but mortality is accelerated in both groups at higher heat stress temperatures (37-38°C)^22^. Taken together, the results of our study along with those of Jaramillo and Castaneda ^22^ indicate there may be a window of temperatures at which gut microbial communities can enhance host heat tolerance, but there is an upper limit of temperatures at which animals succumb to heat stress regardless of microbial colonization. Interestingly, the flies used by Jaramillo and Castaneda ^22^ were completely devoid of bacteria, while we were able to obtain similar results using tadpoles that were still colonized with bacterial communities, but had depleted diversity in these communities. Our results are also similar to those that found an association between losses of gut bacterial diversity and reduced animal survival in lizards^27^. However, the lizard study was correlative and thus could not disentangle the direct impact of microbial diversity from other physiological impacts that may affect both bacterial diversity and host survival.

We have thus shown for the first time that directly manipulating the host-associated microbiome can impact animal survival under heat stress in an ectothermic vertebrate. These findings are important because the temperatures tested are consistent with heat wave air temperatures in our study area (PA, USA), that are expected to become increasingly common in future decades^76^. Further, anthropogenic impacts and increasing temperatures are resulting in a widespread loss of host-associated microbial diversity^54^, and therefore, these effects could synergize to produce negative impacts on ectotherm survival in the wild.

### Experimental depletion of tadpole microbiota altered the thermal sensitivity of locomotor performance

To determine if the impacts of gut microbiota that we observed on tadpole acute thermal tolerance and fitness would also affect the thermal sensitivity of whole-organism performance, we measured locomotion performance of tadpoles across a range of temperatures. Specifically, after the completion of the survival portion of experiment 2, we measured the maximum swimming velocity in 18 colonized and 18 depleted tadpoles from the 22°C control group at six assay temperatures: 5, 15, 22, 26, 30, and 34°C. The velocity measures were then standardized to the body length of each individual to control for body size differences across groups.

As predicted, temperature significantly impacted tadpole locomotor performance (Fig. 3; GLMM, χ^2^= 242.01, p<0.001), such that performance tended to increase with increasing temperature. Locomotor performance typically follows the standard thermal performance curve relationship where performance increases steadily with temperature until reaching an optimum and then declines sharply^77, 78^. However, our data indicates that we did not reach temperatures at which this decline is observed and thus we were unable to calculate the thermal optimum. This may be because we measured up to only 34°C, which is still several degrees below our observed values of CT_max_ (∼40°C), and locomotor performance has been observed to increase until close to the upper lethal limits^78^.

**Fig. 3.**
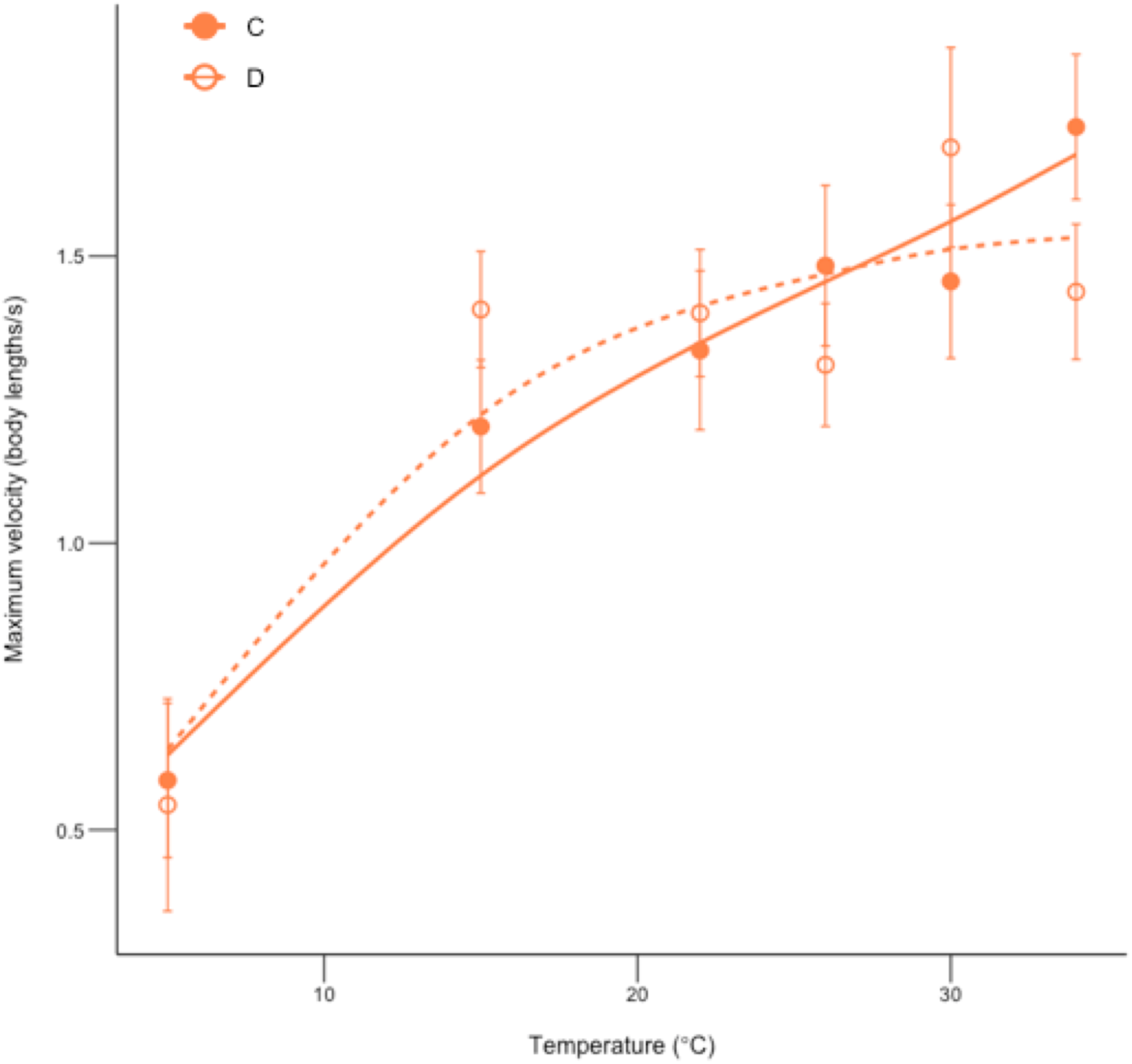
Maximum swimming velocity of colonized and depleted tadpoles at six assay temperatures. C = colonized tadpoles and D = depleted tadpoles. Error bars show means ± s.e.m. and curves were fit to each group using a GAM function. N = 17-18 animals per group.

Nevertheless, we still observed an effect of microbial colonization treatment on the thermal sensitivity of locomotor performance. Specifically, we found a significant interaction between temperature and microbial colonization treatment on tadpole maximum velocity (Fig. 3; GLMM, χ^2^= 19.89, p=0.02), although we did not observe a direct effect of microbial colonization (GLMM, p>0.05). This interactive effect represents a crossing of colonized and depleted curves at higher assay temperatures (Fig. 3), such that the performance of depleted tadpoles plateaued at higher temperatures, whereas performance continued to increase with temperature in colonized tadpoles.

This result suggests that not only does depleting the gut microbiome result in reduced thermal tolerance and fitness of tadpoles, but depleted tadpoles may also experience a detriment to locomotor performance at high temperatures compared to colonized tadpoles. In addition to the direct effects of heat on tadpole fitness observed previously, reduced locomotor capabilities may further reduce depleted tadpole fitness under heat stress. Locomotor performance is thought to be strongly linked to animal fitness due to its importance in predator escape, prey capture, and reproduction^79, 80^. As tadpoles are mainly herbivores and are non-reproductive, predator escape is likely the most important factor during the larval stage. However, it would be interesting to test if these effects persist in adult frogs after metamorphosis. Although infection status has been shown to impact the shape of thermal performance curves in bacteria^14^, to our knowledge, our study represents the first evidence of an impact of the host associated-microbial community on the shape of a thermal performance curve in an ectothermic animal system.

### Experimental depletion of tadpole microbiota resulted in subtle changes to tail muscle phospholipid composition

After investigating thermal tolerance, fitness, and whole-organismal performance, we tested for differences between colonized and depleted tadpoles in several physiological metrics that may be associated with thermal tolerance to understand how the gut microbiome may impact thermal physiology at lower levels of biological organization, and what factors might underlie our results. The saturation of phospholipids in cell membranes and resulting changes in membrane fluidity is one biological mechanism that can contribute to thermal acclimation in ectotherms^6, 35, 36, 81–83^. Our study species, the green frog tadpole, has been previously shown to modulate phospholipid composition of cell membranes in response to temperature acclimation^36^. For example, winter acclimated tadpoles have higher proportions of unsaturated fatty acids in their muscles than summer acclimated tadpoles^36^. To probe if cell membrane composition may contribute to the differences in thermal phenotypes we observed between colonized and depleted tadpoles, we used LC-MS analysis to compare the composition of phospholipids in tail muscle tissue between colonized and depleted tadpoles from our first experiment, from a single acclimation temperature of 22°C.

We detected twelve phospholipid species which had molar percentages that were significantly different between colonized and depleted tadpoles (Supplementary Table 4; ANOVA, corrected p<0.05). However, in multivariate analyses using Bray-Curtis distance between samples, we did not detect significant changes in the overall phospholipid species community composition across groups (PERMANOVA, p>0.05). We also did not detect significant differences in the molar percentages of any specific phospholipid classes (Phosphatidylcholines (PC), Phosphatidylethanolamines (PE), Phosphatidylglycerols (PG), Phosphatidylinositols (PI), and Phosphatidylserines (PS)) across groups (ANOVA, corrected p>0.05). Lastly, there were no significant differences in the proportions of saturated fatty acids, monounsaturated fatty acids, and polyunsaturated fatty acids, or overall unsaturation indices, peroxidation indices, and average carbon chain length between colonized and depleted tadpoles (two-sided t-tests, p>0.05). In tadpoles and newts, increases or decreases in the latter three metrics have been associated with enhanced acclimation to cold or heat, respectively^36, 81^.

In mouse systems, there are differences in tissue and serum lipid profiles, including abundances of specific phospholipids, between germ-free and conventional animals^84^, which could be due to the role of gut microbiota in phospholipid metabolism^85^. Our results demonstrate that in an amphibian system, depleting the gut microbiota can also result in alterations to the abundances of specific phospholipid species. However, due to the lack of correlation between gut microbial colonization and traits expected to be associated with thermal tolerance (membrane unsaturation index etc.), these differences are likely not the cause of the altered thermal tolerance we observed across groups.

### Experimental depletion of tadpole microbiota reduced mitochondrial enzyme activity

The function of enzymes is another important subcellular mechanism impacting thermal performance in ectothermic animals^6, 35^. For example, the loss of enzyme structure and function at high temperatures can underlie thermal tolerance if it occurs at temperatures lower than those impacting function at higher levels of organization^35^. According to the oxygen capacity limitation of thermal tolerance hypothesis (OCLTT), mitochondrial enzymes, which contribute to oxidative capacity, may be particularly important in maintaining thermal tolerance. This hypothesis posits that thermal tolerance limits at extreme temperatures are set by a mismatch in the supply and demand for oxygen, and a resulting loss of aerobic scope, which is the difference between routine and maximal rates of oxygen consumption^34, 86^. Although there is debate and mixed support surrounding this hypothesis^87, 88^, the activity of aerobic mitochondrial enzymes are positively associated with enhanced thermal performance in both heat and cold in many ectothermic vertebrate systems including several fish^89–93^, alligators^94^, newts^95^, as well as green frog tadpoles^36^.

To evaluate the importance of mitochondrial enzymes to thermal tolerance in our system, we tested the activity of two enzymes, citrate synthase (CS) and cytochrome c oxidase (CCO), in tail muscle tissue from colonized and depleted tadpoles from our first experiment in the 28°C acclimation group. The tissue mass-specific activity of each enzyme was tested at three assay temperatures, 28°C which corresponds to the animal’s acclimation temperature, 34°C which is a temperature at which we observed differential survival between colonized and depleted tadpoles, and 40°C which is close to the CT_max_ values for these animals.

We found that assay temperature had a significant impact on both CS and CCO activities (Fig. 4a-b; repeated-measures ANOVA, F= 76.14 for CCO and 69.07 for CS, p<0.001 for both). For CCO, we found a peak in activity at 34°C, and a significant reduction in activity at 40°C. However, for CS, we found a lower activity at 28°C, with similar levels of activity at 34 and 40°C. The maintenance of CS activity at temperatures close to animal CT_max_ is consistent with the idea that some enzymes are able to maintain function beyond the whole-organism’s critical thermal limit^35^. Regardless of the impacts of assay temperature, we found that depleted tadpoles exhibited significantly lower activities of both CCO and CS activity at all temperatures tested, compared to colonized tadpoles, while controlling for differences in body mass (Fig. 4a-b; repeated-measures ANOVA, F= 5.22 for CCO and 8.75 for CS, p<0.05 for both). Specifically, depleted tadpoles had enzyme activities that were on average 15 and 37% lower than those of colonized tadpoles for CCO and CS respectively.

**Fig. 4.**
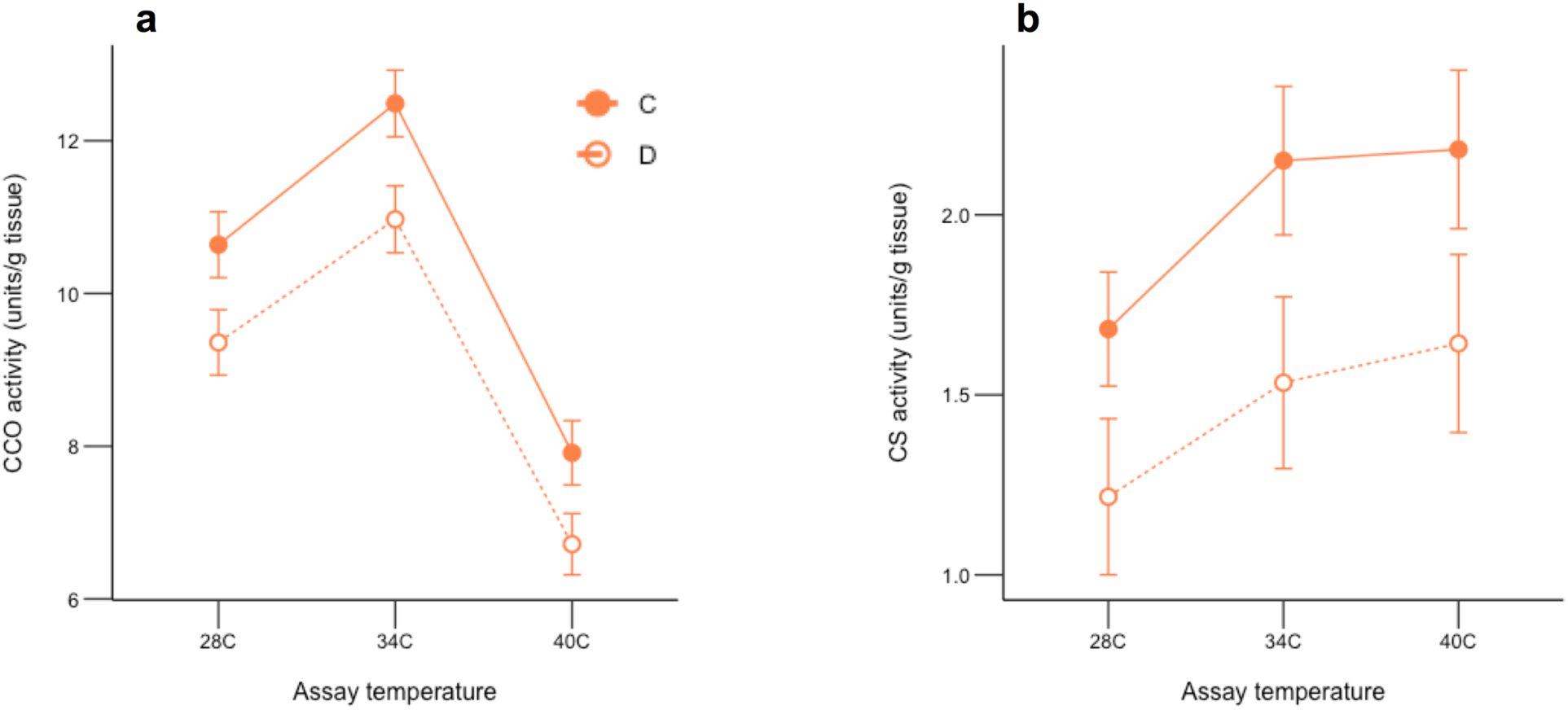
Mitochondrial enzyme activities in tail muscle of colonized and depleted tadpoles at three assay temperatures. **a,** cytochrome c oxidase (CCO) activity **b,** citrate synthase (CS) activity. C=colonized tadpoles and D=depleted tadpoles. Error bars show means ± s.e.m. On the y-axes, one unit is equal to one μmol of substrate modified per minute. N = 7-8 samples per group for each enzyme.

Reductions in mitochondrial enzyme activities in depleted tadpoles could be associated with the lower thermal tolerance we also observed in this group. Activity of CS is positively correlated with mitochondrial density and VO_2_ max^96^, which is the upper limit of aerobic scope, and CCO is thought to be a major regulator of oxidative phosphorylation capacity and ATP production^97^. In accordance with the OCLTT hypothesis^86^, depletion of gut microbiota could reduce tadpole aerobic scope, via reductions in mitochondrial enzyme activity and associated VO_2_ max, and thus limit thermal tolerance in these animals. There are several possible mechanisms by which microbiome depletion may impact mitochondrial phenotypes. Gut microbiota can have marked impacts on mitochondrial metabolism mainly through the action of short chain fatty acids (SCFAs) produced by microbial fermentation. For example, there is a 70% drop in oxidative phosphorylation rates in colonocytes of germ-free mice, which can be rescued by the addition of butyrate, a SCFA serving as the primary energy source for cells in the colon^98^. In a congeneric species of the green frog, the American bullfrog, an estimated 20% of daily energy requirements during the tadpole stage come from microbially-produced SCFAs in the gut^99^. Moreover, production of SCFAs and bile acids by gut microbiota can stimulate mitochondrial biogenesis and protect against reactive oxygen species (ROS), which allows for an increase in maximal oxygen consumption by hosts during periods of high demand^100^. Future studies measuring levels of SCFAs in the gut contents of colonized and depleted tadpoles are needed to determine if depleted individuals exhibit a loss of mitochondrial enzyme activity due to reductions of important microbial products.

### Interactions between environmental temperature, microbial colonization state, and body mass influenced whole-animal resting metabolic rate

Since our mitochondrial enzyme results suggested that microbial depletion could limit tadpole aerobic scope, and thus thermal tolerance, we explored this idea further by measuring whole-organism resting metabolic rate, which is the lower limit of aerobic scope. A fundamental prediction of the OCLTT hypothesis is that aerobic scope is lost at extreme temperatures when resting metabolic rate meets or exceeds maximal metabolic rate^86^. For heat tolerance, this prediction is based on the idea that routine oxygen consumption tends to increase steeply with rising temperatures, until demand can no longer be met^86, 101–103^. While we did not explicitly measure aerobic scope, we tested the impacts of heat on routine oxygen consumption. Specifically, with the animals remaining in our second experiment, we measured the resting mass-specific metabolic rate (VO_2_) of 10 colonized and 10 depleted tadpoles each at two temperatures: 22°C or after a 24- hour exposure to 32°C.

We found that mass-specific metabolic rate was influenced by temperature treatment, such that animals at the warmer temperature had significantly higher resting metabolic rates compared to those at the cooler temperature (Supplementary Figure 7; GLM, F= 5.63, p<0.001). This result was expected, as positive relationships between temperature and metabolism are a generality across ectothermic organisms^104^. However, regardless of temperature treatment, we did not see a direct effect of microbial colonization treatment on tadpole mass-specific metabolic rate (Supplementary Figure 7; GLM, p>0.05). This result is in contrast to a previous study which found that raising tadpoles in sterile water did alter metabolic rate, although the direction of this effect was dependent on the developmental stage of the animals^105^.

However, we did find that there was a significant effect of the interaction between microbial colonization state and body mass on mass-specific metabolic rate, that also depended on assay temperature. Specifically, we saw that at the cooler assay temperature (22°C), both colonized and depleted animals exhibited a significant negative relationship between mass-specific VO_2_ and body mass (Fig. 5a; GLM, F= 5.99, p=0.02). This relationship is consistent with the theory of metabolic scaling^106^, which explains that across a range of animal taxa, larger animals tend to have higher absolute metabolic rates, but lower mass-specific metabolic rates, because mass and whole-animal metabolic rates do not scale linearly. At the higher assay temperature (32°C), there was still a strong negative relationship between mass-specific VO_2_ and body mass in colonized tadpoles, but no relationship between these variables in depleted animals (Fig. 5b; GLM, mass x treatment interaction, F= 5.48, p=0.03).

**Fig. 5.**
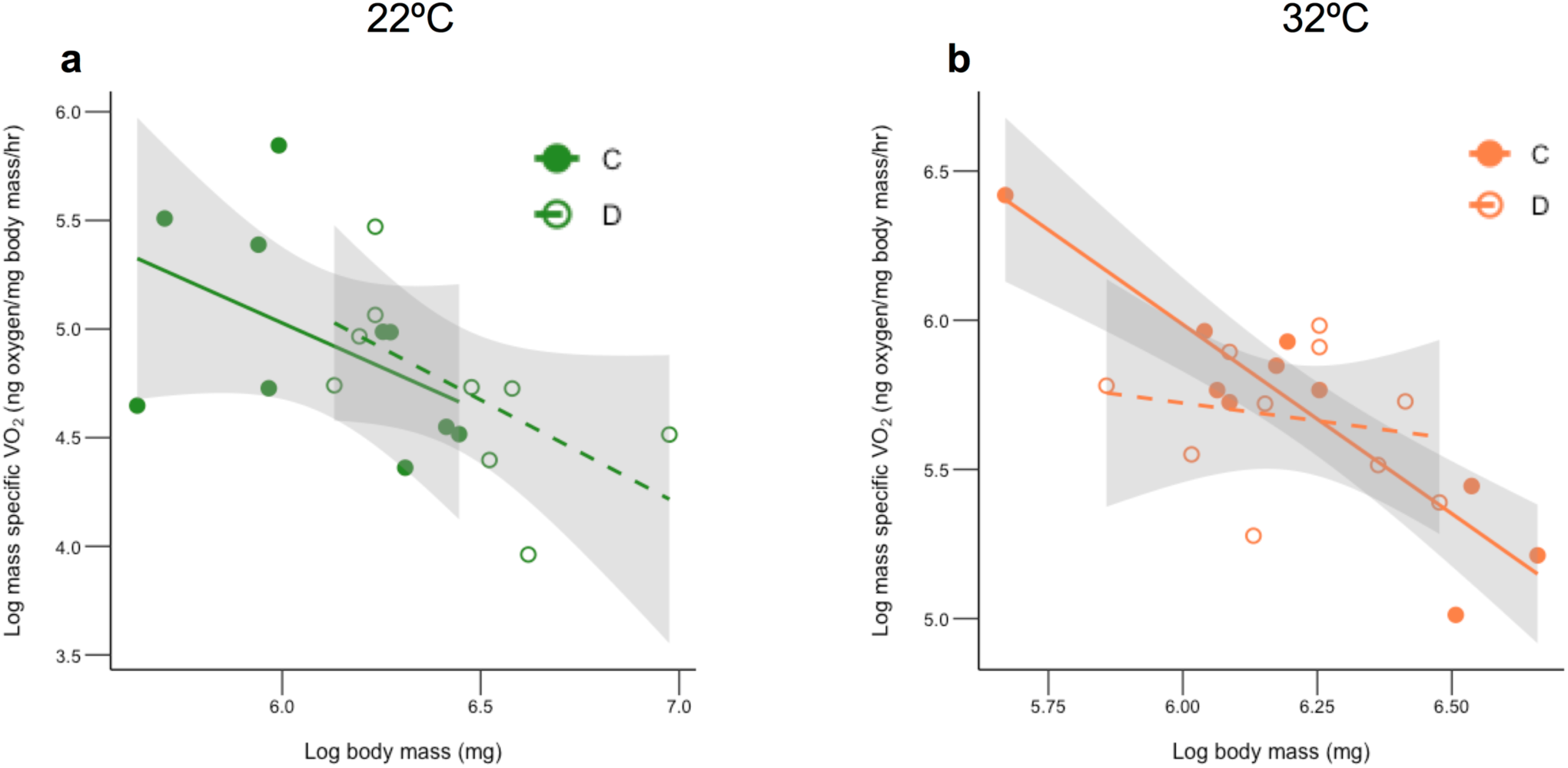
Relationship between mass-specific resting metabolic rate and body mass in colonized and depleted tadpoles at two temperatures. **a,** results from assays performed at 22°C **b,** results from assays performed at 32°C. C=colonized tadpoles and D=depleted tadpoles. Gray shading indicates 95% percent confidence intervals. On the y-axes VO_2_ refers to oxygen consumption.

This result suggests that larger depleted tadpoles may have higher mass-specific metabolic rates than colonized individuals of the same size. It is possible this increase in metabolic rate in depleted animals is due to the interaction of several stressors (high temperature and microbial depletion). Physiological responses to stress can be energy intensive and exposure to several stressors can often result in increases in resting metabolic rates in ectotherms^107–109^. These differences may occur only in larger animals because some studies show that temperature stress may occur more quickly in larger bodied ectotherms^110^. Regardless of the mechanisms, an increase in oxygen demand in large depleted animals at high temperatures coupled with a lower oxidative capacity of the mitochondria in these individuals, as evidenced above, can be expected to result in a narrower window of aerobic scope at high temperatures. Ultimately, these effects may result in the reduced heat tolerance we observed in deleted tadpoles as compared to colonized tadpoles. Although this hypothesis is promising, an explicit test of aerobic scope across a range of temperatures in colonized and depleted tadpoles is necessary to support it further.

## Conclusion

We have shown for the first time that experimentally reducing the diversity of the gut microbiome results in reduced host thermal tolerance, impaired fitness under heat stress, and an altered thermal sensitivity of whole-organism performance in a vertebrate ectotherm. Increased temperatures themselves may also deplete the gut microbiome, and thus climate warming occurring in wild habitats may further reduce the ability of organisms to cope with increasing temperatures. We additionally present preliminary data assessing the putative mechanisms behind these impacts and suggest a role for microbial depletion-induced changes in mitochondrial performance and metabolism in setting thermal tolerance limits. Future experiments may focus on comparing SCFA production and aerobic scope across groups, as well as assessing any potential global changes in gene expression to further explore the mechanistic basis of these effects. Nevertheless, our results coupled with those from invertebrate studies, suggest gut microbial communities are an important factor impacting the response of animals to environmental temperature, and should be considered when using mechanistic models based on individual-level differences in physiology to predict species’ responses to global climate change^111^.

## Methods

### Animal and water collections

All animal research was approved by the University of Pittsburgh IACUC under protocol no. 18062782. Adult green frogs (*Lithobates clamitans*) (5 males, 5 females) used in experiment 1 were collected from a single ephemeral pond in May 2019 from Kisatchie National Forest (LA, USA) and shipped overnight to the University of Pittsburgh. Adult green frogs (5 males, 5 females) used in experiment 2 were collected in July 2020 from a single pond at Pymatuning Laboratory of Ecology (PA, USA) and transported via automobile to the University of Pittsburgh. Permission to collect animals was obtained from LA Department of Wildlife and Fisheries (Scientific Collecting Permit WDP-19-010) and PA Fish and Boat Commission (Scientific Collector’s Permit 2020-01-0131). In the laboratory, frogs were housed individually in a climate-controlled animal room maintained at 24°C, 65% humidity, and on a 14hr: 10hr light: dark cycle prior to experimentation. In June 2019, we collected 100L (divided into 10L carboys), and two smaller (1mL each) samples, of pond water from same location in Louisiana where frogs were collected. In 2020, 100L of pond water was collected monthly from July-October from the same location where frogs were collected in Pennsylvania. The carboys were kept cold on ice, and the 1mL samples frozen on dry ice, and driven back to the University of Pittsburgh. Pond water was filtered through a 500μm sieve to remove macroorganisms and stored at 4°C until use. The 1mL samples were frozen at -80°C.

### Adult spawning and tadpole microbial colonization manipulation

To obtain tadpoles for experimentation, in September 2019 (for experiment 1) and July 2020 (for experiment 2), we induced spawning in a single pair of adult frogs using a hormonal injection method as described by Trudeau, et al. ^112^. Specifically, frogs were injected intraperitoneally with a hormonal cocktail containing GnRH-A (0.4 μg/g body weight, Sigma Aldrich), metoclopramide (10 μg/g body weight, Sigma Aldrich), 0.7% saline (4 μL/g body weight) and DMSO (1 μL/g body weight) at a concentration of 7.5μL x body weight. Mating pairs were then placed in 55-gallon polypropylene tubs filled with ∼4 inches of laboratory water for spawning. Tubs contained pieces of floating styrofoam and PVC half pipes to be use as retreats. Spawning tubs were checked daily for up to one week for egg production. After egg mass production, viable embryos were separated from unfertilized eggs and placed in a 16-quart polypropylene container filled halfway with autoclaved laboratory water, which was changed daily to provide sufficient oxygenation to embryos. After hatching and development to Gosner stage^56^ 25, tadpoles were divided into two treatment groups. Tadpoles assigned to a “colonized” treatment group were placed in a water treatment consisting of 75% autoclaved laboratory water and 25% unmanipulated pond water from their parent’s site of capture to provide a natural source of gut microbiota. Tadpoles assigned to a “depleted” treatment group were placed in a water treatment consisting of 75% autoclaved laboratory water and 25% autoclaved pond water to remove the primary source of gut microbiota^38^. A single clutch of eggs was used for each experiment to reduce the genetic variation in our study subjects.

### Experiment 1 design

We distributed 180 tadpoles spawned from frogs collected from Louisiana into 36 1L polypropylene containers (five individuals per container). Colonized tadpoles were assigned to half of the containers which were filled with autoclaved laboratory water (675 mL), seeded with unmanipulated pond water collected from the parent’s site of capture (225 mL). Depleted tadpoles were assigned to the other half of the containers which were filled with autoclaved laboratory water (675 mL), seeded with autoclaved pond water (225 mL). We placed 12 containers (six colonized and six depleted) each into three 8.5-gallon polycarbonate water baths. The animal chamber was set to maintain water temperature at 14°C (the lowest acclimation temperature) and kept at 65% humidity on a 14hr: 10hr light: dark cycle. Water temperature in all three baths was kept at 22°C using aquarium heaters. All tadpoles were allowed to develop at this temperature for four weeks. We then increased temperature in one water bath to 28°C (using aquarium heaters), and decreased temperature in one water bath to 14°C (by removing aquarium heaters). The third water bath remained at 22°C and animals were allowed to acclimate for another three weeks.

Throughout development, we fed each container of tadpoles weekly a diet of three 0.5g blocks of autoclaved ground rabbit chow suspended in autoclaved agar and supplemented with a small amount (∼1 tsp) of commercial pet vitamins (Reptivite, Zoo Med Laboratories). Tadpoles were transferred weekly to fresh containers filled with the appropriate water treatment. Each week we collected a 1mL sample of stored pond water during water changes to monitor changes in pond water microbial communities through storage. These samples were frozen at -80°C after collection. We monitored water temperature in each tadpole container daily, adjusting aquarium heater temperatures if needed to stay within 1°C of our target treatment temperatures. Ultimately, mean (± sd) temperatures across all tanks were 14.4 ± 0.1, 22.0 ± 0.1, and 28.0 ± 0.3 for our cool, moderate, and warm temperature treatments respectively.

### Acute thermal tolerance assays and animal dissections

After the acclimation period, we tested tadpole acute thermal tolerance by measuring the critical thermal minimum (CT_min_) and maximum (CT_max_) following Hutchison’s dynamic method^55^. Specifically, individual tadpoles were placed in 400mL beakers that contained 200mL of room temperature laboratory water. Beakers were then submerged halfway in a refrigerated circulating water bath (Arctic A10B, Fisher Scientific). To measure CT_min_ and CT_max_, the temperature of the bath was decreased or increased by 0.5°C per minute, respectively. Every minute each tadpole was prodded with a small metal spatula until it no longer responded to the stimulus and could not right itself. We then measured the temperature of the water immediately adjacent to the tadpole using a Traceable Type K thermometer (Fisher Scientific) and placed the animal in another beaker of room temperature water for recovery. Animals were monitored for at least one hour prior to euthanasia and all animals successfully recovered. Individual tadpoles only underwent one trial (CT_min_ or CT_max_) and we conducted 11 trials for both CT_max_ and CT_min_, during which one animal from each of the six treatment groups was measured (three acclimation temperatures x two microbial colonization treatments). The remaining individuals from the experiment were euthanized and dissected without collecting thermal tolerance data.

Each individual was euthanized by immersion in buffered MS-222 (10g/L). Following euthanasia, we recorded tadpole body mass (g), and placed each individual under a Leica S9i dissecting microscope fitted with a camera attachment to photograph each tadpole and record their Gosner stage. Photographs were later used to measure tadpole body length and width (mm), excluding the tail. We also measured facial symmetry by measuring the distance (mm) from the center of each eye to the tip of the nose and computing the absolute value, subtracted from 1, of the difference between these two measures. All images were analyzed using Image J v1.52q. We then dissected each individual and removed the entire gastrointestinal tract for microbiome and flow cytometry analyses, and the entire tail for mitochondrial enzyme and phospholipid membrane analyses. Gut samples were immediately frozen at -80°C and tail samples were flushed in N_2_ gas, frozen under liquid N_2_, and then stored at -80°C. Dissection instruments were wiped with 70% EtOH and flame sterilized between each individual.

To assess the effect of acclimation temperature, microbial colonization treatment, and their interaction on tadpole CT_min_, CT_max_, body mass, body width, body length, Gosner stage, and facial symmetry, we used generalized linear mixed models (GLMMs) including tadpole tank as a random effect in all models. Body length was included as a covariate in the facial symmetry model, and body mass and Gosner stage were included as covariates in the CT_min_ and CT_max_ models. All models were constructed in R v3.4.3^113^ using the lme4 package^114^.

### Flow cytometry

We used flow cytometry to calculate the absolute bacterial abundance in gut samples from experiment 1. Whole gut samples were split in half and one half was used for microbiome analyses (see below), and the other was used for flow cytometry. We performed this analysis on 9-11 samples per treatment group, except for the 14°C colonized group. In this group, we only retained four samples as most were too small to be divided and we prioritized microbiome analyses. For each sample, gut contents were removed from the gut tissue and weighed. Specifically, the gut was unraveled on a weigh boat and divided into short sections with a sterile spatula and inoculating loop. For each section, the contents were lightly scraped out of the tissue. After all contents were removed, the tissue was discarded, and the weigh boat was flushed with 1mL sterile PBS. The gut contents and the PBS were carefully transferred to a pre-weighed microcentrifuge tube which was spun at 12,000 rpm for 10 minutes. The PBS was then pipetted off and the contents weighed. The gut content samples were then diluted in sterile PBS at a ratio of 100mL PBS to 1g of gut contents, and the resulting suspension was filtered through a 5μm cell strainer. To create a stock solution, 949μL of the cell suspension was mixed with 50μl CountBright Absolute Counting Beads (Thermo Fisher Scientific) as an internal reference, and 3μL of a 1:100 dilution of SYBR Green I dye (Sigma Aldrich). We then diluted this stock solution 1:4 with PBS.

We counted bacterial cells on the Attune NxT Flow Cytometer (Thermo Fisher Scientific). We initially adjusted the gating of the machine to count populations of both counting beads and stained bacterial cells. Specifically, plots of fluorescein isothiocyanate (FITC) were used to distinguish stained cells from all other events, and forward scatter (FSC) vs. side scatter (SSC) plots were used to distinguish counting beads from all other events and identify single prokaryotic cell events (Supplementary Figure 8). To establish initial gates for counting beads, blank samples spiked only with beads were used. We ran 100μL of each sample on the machine and counted beads and bacterial cells until 500,000 events were reached, with a flow rate of 12.5 μL/min, and a threshold of 0.3. Instrument and gating parameters were kept constant across all samples. Based on the number of beads and bacterial cells counted, we then calculated the number of cells/g of gut contents in each sample following the manufacturer’s protocol. To compare the abundance of bacterial cells across treatment groups, we used a log transformation to normalize the cell counts, and used a linear mixed effect model in the nlme^115^ package in R with acclimation temperature, microbial colonization treatment, and their interaction as fixed effects, and the date of measure as a random effect. We removed two outliers from analysis (n=1 from the 14°C colonized group and n=1 from the 28°C colonized group).

### Phospholipid analyses

We identified the phospholipids present in tail muscle tissue of eight colonized and eight depleted tadpoles from the 22°C group only from experiment 1. Tail tissue samples were sent to the University of Pittsburgh’s Health Sciences Metabolomics and Lipidomics core facility for LC-MS analysis to identify the phospholipids present in each sample. At the facility, muscle samples were prepared for metabolic quenching, lysis, and lipid extraction by adding 500µL ice cold PBS and were homogenized using MP Bio Matrix A tubes at 60hz for 1 minute. 400µL of uncleared supernatant was transferred to a clean glass tube containing 10µL each PI, PC, PS, PE, and PG UltimateSplash deuterated phospholipid internal standards (Avanti Polar Lipids) and subjected to a Folch extraction. Samples were rested on ice for 10 minutes before phase separation via centrifugation at 2500 x g for 15 minutes. 700µL of organic phase was dried under nitrogen gas and resuspended in 1:1 acetonitrile:isopropanol and 3µL of sample was subjected to online LC-MS analysis.

Analyses were performed by untargeted LC-HRMS. Briefly, samples were injected via a Thermo Vanquish UHPLC and separated over a reversed phase Thermo Accucore C-18 column (2.1×100mm, 5μm particle size) maintained at 55°C. For the 30 minute LC gradient, the mobile phase consisted of the following: solvent A (50:50 H2O:ACN 10mM ammonium acetate / 0.1% acetic acid) and solvent B (90:10 IPA:ACN 10mM ammonium acetate / 0.1% acetic acid). Initial loading condition is 30% B. The gradient was the following: Over 2 minutes, increase to 43%B, continue increasing to 55%B over 0.1 minutes, continue increasing to 65%B over 10 minutes, continue increasing to 85%B over 6 minutes, and finally increasing to 100% over 2 minutes. Hold at 100% for 5 minutes, followed by equilibration at 30%B for 5 minutes. The Thermo IDX tribrid mass spectrometer was operated in both positive and negative ESI mode. A data-dependent MS2 method scanning in Full MS mode from 200 to 1500 m/z at 120,000 resolution with an AGC target of 5e4 for triggering ms2 fragmentation using stepped HCD collision energies at 20,40, and 60% in the orbitrap at 15,000 resolution. Source ionization settings were 3.5 kV and 2.4kV spray voltage respectively for positive and negative mode. Source gas parameters were 35 sheath gas, 5 auxiliary gas at 300°C, and 1 sweep gas. Calibration was performed prior to analysis using the PierceTM FlexMix Ion Calibration Solutions (Thermo Fisher Scientific). Internal standard peak areas were then extracted manually using Quan Browser (Thermo Fisher Xcalibur ver. 2.7) and normalized to weight and internal standard peak area.

Using this data, each phospholipid species present in each sample was identified by class (PC, PE, PG, PI, or PS) and species were grouped together based on their class, their total carbon chain length, and total number of double bonds (combined for both fatty acid tails). We calculated the molar percentage of each phospholipid class, and each phospholipid species out of the total for each sample. We additionally calculated the combined molar percentages of all saturated fatty acids (%SFA; zero double bonds), monounsaturated fatty acids (%MUFA; single double bond), and polyunsaturated fatty acids (%PUFA; >1 double bond) for each sample. We then calculated the unsaturation and peroxidation indices for each sample as described by Hulbert, et al. ^116^. The unsaturation index computes the number of double bonds per 100 fatty acids, and the peroxidation index determines the susceptibility of membranes to peroxidation. Lastly, we calculated the average chain length for each sample by dividing the molar percentage of each phospholipid species in the sample by its chain length, summing these values, and dividing by 100.

To identify differences between colonization treatment groups in the molar percentages of individual phospholipid classes or species, we used the response screening function in JMP v14.1 which performs ANOVAs across groups with the abundance of each phospholipid class or species as the response variable and subsequently corrects all p-values using the FDR method. For phospholipid species, we included only those that were present at greater than 0.1% relative abundance in at least one treatment group. To compare phospholipid compositions across colonization groups in multivariate space, we created a Bray-Curtis distance matrix using the vegdist function in the vegan package^117^ in R, based off of our molar percentage data of each phospholipid species, excluding species present in abundances of less than 0.1% in both treatment groups. We then used the adonis2 function to perform a PERMANOVA on the distance matrix, with 999 permutations. Lastly, after confirming normality with Shapiro-Wilk tests, we used two-sided t-tests in JMP to compare %SFA, %MUFA, %PUFA, unsaturation indices, peroxidation indices, and average chain lengths, between the two colonization groups.

### Mitochondrial enzyme assays

We measured the activities of cytochrome c oxidase (CCO) and citrate synthase (CS) isolated from the tail muscle of eight depleted and eight colonized samples from the 28°C acclimation group from experiment 1. Enzyme activities were measured at three assay temperatures: 28, 34, and 40°C following previously published protocols^36^. Some samples were too small to be analyzed individually and, in those cases, we pooled samples from 2-3 individuals together from the same treatment group. To prepare homogenates for enzyme activity assays, frozen tail muscle was homogenized in 9 volumes of homogenization buffer (50 mM imidazole, 2 mM MgCL2, 5 mM EDTA, 1 mM glutathione, and 0.1% Triton X 100, pH=7.5) and centrifuged at 300 g for 10 minutes at 4°C. The supernatant that resulted from centrifugation was used for enzyme assays. The activities of CS and CCO were calculated from the linear change in absorbance with a temperature-controlled spectrophotometer (Evolution 201, Thermo Fisher Scientific) outfitted with an 8-cell changer that was connected to a circulating water bath (1160S, VWR). The activity of CS was determined from the reduction of DTNB (5,5’ dithiobis-2- nitrobenzoic acid) at 412 nm in assay medium containing 100 mM Tris-HCL (pH=8.0), 0.1 mM DTNB, 0.15 mM acetyl-CoA, 0.15 mM oxaloacetate. The activity of CCO was determined from the oxidation of reduced cytochrome c at 550 nm against a reference of 0.075 mM cytochrome C oxidized with 0.33% potassium ferricyanide in assay medium containing 100 mM potassium phosphate (pH=7.5) and 0.075 mM reduced cytochrome c. Cytochrome c was reduced with sodium hydrosulfite, and excess sodium hydrosulfite was removed by bubbling with air for two hours^118^. All assays were performed in duplicate, and enzyme activity is expressed as units per g of tissue, where one unit is equal to 1 µmol of substrate modified per minute.

To determine the effects of both assay temperature and microbial colonization treatment on CS and CCO activity, we used repeated-measures ANOVAs in R, controlling for the effects of individual. In both models, body mass was included as a covariate, and if samples were pooled, the average body mass of the pooled individuals was used instead. In our CS model, we removed one depleted outlier sample from analysis.

### Experiment 2 design

We distributed 200 tadpoles spawned from frogs collected from Pennsylvania individually into 12oz polypropylene containers. Half of the tadpoles were assigned to a colonized treatment, and their containers were filled with autoclaved laboratory water (225mL) and seeded with unmanipulated pond water from the parent’s site of capture (75mL). The other half of the tadpoles were assigned to a depleted treatment and their containers were filled with autoclaved laboratory water (225mL) and seeded with autoclaved pond water (75 mL). The animal chamber was set to 22°C and kept at 65% humidity on a 14hr: 10hr light: dark cycle. Throughout development, we fed the tadpoles weekly a diet of one 0.25g block of autoclaved ground rabbit chow suspended in autoclaved agar, supplemented with a small amount (∼1 tsp) of commercial pet vitamins (Reptivite, Zoo Med Laboratories). Tadpoles were provided fresh water once weekly by transfer to new containers filled with the appropriate water treatment.

After seven weeks of development, we removed 20 tadpoles per treatment and assayed their CT_ma**x**_, following the protocol described above, to verify repeatability of our heat tolerance results from experiment 1 in a second population of tadpoles. After measuring CT_max_, we euthanized tadpoles in buffered MS-222 (10g/L), and recorded their mass (g), body length excluding tail (mm) using digital calipers, and Gosner stage. We then dissected each animal and removed the entire gastrointestinal tract, to be used later for microbiome analyses. Dissection instruments were wiped with 70% EtOH and flame sterilized between each individual. All gut samples were stored at -80°C prior to processing. We used generalized linear models (GLMs) in R to compare body mass, body length, Gosner stage, and CT_max_ between colonized and depleted tadpoles.

We then moved 78 tadpoles (39 from each colonization treatment) into a separate animal chamber kept at 32°C, 65% humidity, and on a 14hr: 10hr light: dark cycle. An additional 78 tadpoles (39 from each colonization treatment) remained in the previous animal chamber at 22°C. After 10 days, we increased the temperature of the heat treatment to 34°C for a duration of 10 days, and to 36°C for a final 10 days. In the 32-36°C temperature treatment group we observed 100% mortality in both treatment groups by the end of the 30 days. In the 22°C group, we did not observe any mortality in either group over the duration of the 30 days and these animals were used for additional experimentation (see below).

To compare survival of colonized and depleted tadpoles under heat stress, we performed a discrete-time logistic regression analysis in R using the base glm function with a binomial distribution, including temperature (synonymous with each 10-day time period) and microbial colonization treatment as predictor variables. We computed the hazard function of mortality for both colonization groups at each temperature from our model using the formula established by Singer and Willett ^119^. Because no mortality was observed, animals at 22°C were not included in the analysis.

### Resting metabolic rate

Of the 78 remaining tadpoles in the 22°C group, 40 were used to assay resting metabolic rate. We measured metabolic rate in 10 tadpoles per microbial colonization treatment group (20 total) at 22°C. The remaining 10 per colonization treatment group (20 total) were placed at 32°C for 24 hours, prior to measuring metabolic rate at 32°C. Resting metabolic rate was measured via oxygen consumption (VO_2_) using an intermittent respirometry system (Q-box mini-AQUA aquatic respirometer, Qubit Systems). Prior to measurements, tadpoles were weighed in grams, and their volume calculated via water displacement in a 10mL graduated cylinder. Tadpoles were then placed in the 9mL respirometry chamber and rested for five minutes. We then measured eight cycles of oxygen consumption per animal, each of which consisted of an 87s flush phase, a 15s rest phase, and a 360s closed circulation phase. We exited the room during data collection to minimize stress to tadpoles, though small movements of tadpoles could not be prevented. Immediately following data collection, tadpoles were euthanized via immersion in buffered MS-222 (10 g/L), and their Gosner stage was recorded.

For each of the eight cycles of intermittent respirometry per animal, mass-specific VO_2_ was calculated in ng O_2_/ mg body mass/hour using the formula: O_2_ slope (ng/L/sec) * (respirometer volume (L)-animal volume (L)) * 3600s / body mass (mg). Because we could not account for small movements of animals, to best approximate resting metabolic rate, the minimum value of VO_2_ of the eight measurements per animal was used for analyses. Prior to analyses, VO_2_ values were normalized using a log transformation. We used GLMs in R to test for significant effects of assay temperature, microbial colonization treatment, body mass, and interactions of these variables on mass-specific VO_2_. Gosner stage was included as an additional covariate in the model. Two depleted tadpoles (one from 22°C, and one from 32°C) were removed from analysis because they did not survive the assay.

### Locomotion performance

Of the remaining animals from the 22°C treatment group, 18 per microbial colonization treatment (36 total) were shipped overnight to Elmhurst University where the thermal sensitivity of locomotor performance was measured. Animal work was approved by Elmhurst University IACUAC protocol no. FY21-007. We measured the maximum swimming velocity of colonized and depleted tadpoles at 5, 15, 22, 26, 30 and 34°C in a water-jacketed 70 cm x 10 cm x 5 cm plexiglass track containing 3 cm of laboratory water. The bottom of the track was marked in 1 cm increments and the water jacket of the track was connected to a circulating water bath (1160S, VWR) that controlled the temperature of the water in the track. Prior to locomotor performance assays, the tadpoles were brought from their housing temperature (22°C) to the appropriate assay temperature at a rate of 5°C/hr in a temperature-controlled biological incubator (model I-41 VL, Percival). Tadpoles remained at the assay temperature for 30 min prior to assays. The order of assay temperatures was 22, 5, 30, 15, 34, and 26°C, and velocity was measured at 1 assay temperature per day with 24 hours between assays at different temperatures.

To measure locomotor performance, we prompted tadpoles to swim by touching their tail with a blunt probe and filmed their escape response (frame rate = 60 fps) with a video camera (HC-V180, Panasonic) positioned 90° above the track. Three escape responses were recorded for each tadpole at a particular assay temperature, and each escape response was analyzed frame-by-frame using Tracker Video Motion Analysis and Modeling Tool software v5.1.5 (Open Source Physics,www.opensourcephysics.org) to calculate the maximum velocity (cm/s) reached during the first 10 cm for each escape response. We used the highest velocity measure calculated from the three escape responses for our analysis, and standardized these measures to body length (including tail) so maximum velocity was expressed in units of body lengths/s. Tadpoles that failed to swim at a particular assay temperature were omitted from our analysis at that temperature (n=1 individual from the colonized group at 5°C, n=1 individual from the depleted group at 5°C, n=1 individual from the depleted group at 22°C, n=1 individual from the colonized group at 26°C, n=1 individual from the colonized group at 30°C and n=1 individual from the colonized group at 34°C).

We used GLMMs in the lme4 package in R to determine the effects of assay temperature, microbial colonization treatment, and their interaction on tadpole maximum velocity and included individual as a random effect. Body length was included as an additional covariate in the model. To visualize the relationship between temperature and performance we fit generalized additive models (GAMs) to our data for both colonized and depleted tadpoles using the mgcv package^120^. For GAMs, a k value of 6 was used.

### Microbiome analyses

We conducted bacterial inventories for gut samples from experiments 1 and 2, and water samples from experiment 1. Total DNA was extracted from gut and water samples, along with blank extraction controls (one from experiment 1 and two from experiment 2), with the QIAamp PowerFecal Pro DNA Isolation Kit (QAIGEN) following the manufacturer’s protocol. Water samples were vortexed after thawing and 200μL of water was used to begin the extraction. Extracted DNA was stored at -20°C prior to sequencing. Extracted DNA from experiment 1 was sent to the University of Illinois at Chicago’s DNA Services Facility and extracted DNA from experiment 2 was sent to the University of Connecticut’s Microbial Analysis, Resources, and Services facility for library preparation, PCR, and sequencing. Specifically, the bacterial 16S rRNA gene was amplified using primers 515F and 806R^121^, libraries were pooled together and purified, and amplicons were then sequenced on the Illumina Miseq platform, resulting in 2×250 paired-end reads. Raw sequence data was processed using QIIME2 v2019.7^122^, and DADA2 was used to trim reads to a length of 180 base pairs, filter reads for quality, merge forward and reverse reads, remove chimeric and singleton reads, and assign reads to amplicon sequence variants (ASVs)^123^. Subsequently, sequences were aligned, a phylogenetic tree was built using FastTree^124^, and ASVs were assigned taxonomy using the SILVA database classifier v132^125^. We removed any sequences identified as mitochondria, chloroplast, or archaea. Because kit control samples returned very low read counts compared to experimental samples (<1000 reads in experiment 1, <10 reads in experiment 2), and ASVs found in kit controls can often be found in the gastrointestinal tract^126^, we did not filter out kit control ASVs from experimental samples. However, we extracted samples in mixed batches, so any minor influences of contamination should be distributed equally across treatment groups^127^.

We created several rarefied ASV tables to compare microbial diversity metrics across experimental groups. For experiment 1, we created an ASV table which contained both gut and water samples and was rarefied to 1,567 sequences per sample (the number in the sample with the fewest reads). We then created an ASV table which was also rarefied to 1,567 sequences per sample and contained only water samples. Lastly, we created a table that contained only gut samples and was rarefied to 14,232 sequences per sample (the number in the sample with the fewest reads). For experiment 2 samples, we created an ASV table containing only gut samples that was rarefied to 13,216 sequences per sample which excluded three samples per microbial colonization group from analysis due to low read counts. For each table, we calculated metrics of alpha diversity within each sample (Shannon diversity^128^, Faith’s phylogenetic diversity^129^, Pielou’s evenness^130^, and the number of bacterial ASVs), and beta diversity between each pair of samples (Bray-Curtis dissimilarity^131^, and unweighted and weighted UniFrac distance^132^) within QIIME2.

For experiment 1, we first compared the composition of bacterial communities between gut and water samples using a PERMANOVA with 999 permutations based on the Bray-Curtis distance between samples in QIIME2. We then repeated this process to compare the community composition between water samples that were collected fresh from the pond or stored in the laboratory. Next, focusing on gut samples only, we examined the effects of both acclimation temperature, microbial colonization treatment, or their interaction on tadpole gut bacterial alpha diversity. We used GLMMs in the lme4 package in R, including tank as a random effect, to compare differences in Shannon diversity, Faith’s phylogenetic diversity, Pielou’s evenness, and the number of observed ASVs. To determine differences in community composition of the microbiome across treatment groups, we used the adonis2 function in the vegan package to perform PERMANOVAs, with 999 permutations, using Bray-Curtis, Unweighted UniFrac, and Weighted UniFrac distance matrices based on both microbial colonization treatment, acclimation temperature, and their interaction. We additionally controlled for the effect of tank using the strata function. We used the same three distance matrices to compare intraindividual variability in the bacterial community across acclimation temperature and microbial colonization treatment groups using PERMDISP tests with the betadisper function. We used the MaAsLin2 package^133^ in R to identify bacterial phyla and genera in gut bacterial communities with relative abundances that were correlated with acclimation temperature or microbial colonization treatment. MaAsLin2 was run using an arcsin-square root transformation of the relative abundance data, and p-values were corrected using the FDR method.

For experiment 2, the four metrics of gut bacterial alpha diversity were compared between colonized and depleted tadpoles using Kruskal-Wallis tests in QIIME2. Bacterial community composition was compared between colonized and depleted tadpoles using the three calculated distance matrices with PERMANOVAs, using 999 permutations, in QIIME2.

## Data availability

Raw microbiome sequencing data is available from NCBI under accession PRJNA732310. All other raw data sets used in this study are available from the Zenodo repository at 10.5281/zenodo.4894888.

## Acknowledgements

We thank Drs. Michel Ohmer, Karie Altman, and Emily Hall for field collection assistance, Kendall Kohler, Maya Maurer, Matt Maier, Sarah Reilly, Amanda Haid, Cory Duckworth, and Jaclyn Adams for animal husbandry and DNA extraction assistance, and Dr. Nick Barts for technical assistance. We also thank the University of Pittsburgh’s Health Sciences Metabolomics and Lipidomics core facility (NIH S10OD023402 PI Wendell), the DNA Services Facility at the University of Illinois at Chicago, and the Microbial Analysis, Resources, and Services Facility at the University of Connecticut for sample processing. This work was supported by the University of Pittsburgh (start-up funds to K.D.K), Elmhurst University (faculty research grant to P.M.M) and the National Science Foundation (GRFP to S.S.F).

## Author contributions

S.S.F, K.D.K, and P.M.M designed the study, S.S.F and P.M.M collected data, S.S.F analyzed the data, generated figures and wrote the initial manuscript draft with editing from K.D.K and P.M.M. K.DK supervised the research.

## Competing interests statement

The authors declare no competing interests.

## Notes

### Competing Interest Statement

The authors have declared no competing interest.

## References

1 Paaijmans, K. P. et al. Temperature variation makes ectotherms more sensitive to climate change. Global Change Biology 19, 2373–2380 (2013).

2 Clusella-Trullas, S., Blackburn, T. M. & Chown, S. L. Climatic predictors of temperature performance curve parameters in ectotherms imply complex responses to climate change. The American Naturalist 177, 738–751 (2011).

3 Pounds, J. A. et al. Widespread amphibian extinctions from epidemic disease driven by global warming. Nature 439, 161 (2006).

4 Sinervo, B. et al. Erosion of lizard diversity by climate change and altered thermal niches. Science 328, 894–899 (2010).

5 Pacifici, M. et al. Assessing species vulnerability to climate change. Nature Climate Change 5, 215–224 (2015).

6 Angilletta Jr, M. J. & Angilletta, M. J. Thermal adaptation: a theoretical and empirical synthesis. (Oxford University Press, 2009).

7 Sunday, J. M., Bates, A. E. & Dulvy, N. K. Global analysis of thermal tolerance and latitude in ectotherms. Proceedings of the Royal Society B: Biological Sciences 278, 1823–1830 (2011).

8 Jørgensen, L. B., Malte, H. & Overgaard, J. How to assess *Drosophila* heat tolerance: unifying static and dynamic tolerance assays to predict heat distribution limits. Functional Ecology 33, 629–642 (2019).

9 Perry, G. M., Danzmann, R. G., Ferguson, M. M. & Gibson, J. P. Quantitative trait loci for upper thermal tolerance inoutbred strains of rainbow trout (*Oncorhynchus mykiss*). Heredity 86, 333–341 (2001).

10 Healy, T. M. & Schulte, P. M. Factors affecting plasticity in whole-organism thermal tolerance in common killifish (*Fundulus heteroclitus*). Journal of Comparative Physiology B 182, 49–62 (2012).

11 Hu, X. P. & Appel, A. G. Seasonal variation of critical thermal limits and temperature tolerance in Formosan and eastern subterranean termites (Isoptera: Rhinotermitidae). Environmental Entomology 33, 197–205 (2004).

12 Nyamukondiwa, C. & Terblanche, J. S. Thermal tolerance in adult Mediterranean and Natal fruit flies (*Ceratitis capitata* and *Ceratitis rosa*): effects of age, gender and feeding status. Journal of Thermal Biology 34, 406–414 (2009).

13 Greenspan, S. E. et al. Infection increases vulnerability to climate change via effects on host thermal tolerance. Scientific Reports 7, 1–10 (2017).

14 Padfield, D., Castledine, M. & Buckling, A. Temperature-dependent changes to host– parasite interactions alter the thermal performance of a bacterial host. The ISME Journal 14, 389–398 (2020).

15 Alberdi, A., Aizpurua, O., Bohmann, K., Zepeda-Mendoza, M. L. & Gilbert, M. T. P. Do vertebrate gut metagenomes confer rapid ecological adaptation? Trends in Ecology & Evolution 31, 689–699 (2016).

16 Kohl, K. D. & Carey, H. V. A place for host–microbe symbiosis in the comparative physiologist’s toolbox. Journal of Experimental Biology 219, 3496–3504 (2016).

17 Fontaine, S. S. & Kohl, K. D. Optimal integration between host physiology and functions of the gut microbiome. Philosophical Transactions of the Royal Society B 375, 20190594 (2020).

18 Ziegler, M., Seneca, F. O., Yum, L. K., Palumbi, S. R. & Voolstra, C. R. Bacterial community dynamics are linked to patterns of coral heat tolerance. Nature Communications 8, 14213 (2017).

19 Russell, J. A. & Moran, N. A. Costs and benefits of symbiont infection in aphids: variation among symbionts and across temperatures. Proceedings of the Royal Society B: Biological Sciences 273, 603–610 (2006).

20 Montllor, C. B., Maxmen, A. & Purcell, A. H. Facultative bacterial endosymbionts benefit pea aphids *Acyrthosiphon pisum* under heat stress. Ecological Entomology 27, 189–195 (2002).

21 Herrera, M. et al. Unfamiliar partnerships limit cnidarian holobiont acclimation to warming. Global Change Biology 26, 5539–5553 (2020).

22 Jaramillo, A. & Castaneda, L. E. Gut microbiota of Drosophila subobscura contributes to its heat tolerance but is sensitive to transient thermal stress. Preprint at https://www.biorxiv.org/content/10.1101/2021.01.08.425860v1 (2021).

23 Moghadam, N. N. et al. Strong responses of *Drosophila melanogaster* microbiota to developmental temperature. Fly 12, 1–12 (2018).

24 Fontaine, S. S., Novarro, A. J. & Kohl, K. D. Environmental temperature alters the digestive performance and gut microbiota of a terrestrial amphibian. Journal of Experimental Biology 221, 187559 (2018).

25 Kohl, K. D. & Yahn, J. Effects of environmental temperature on the gut microbial communities of tadpoles. Environmental Microbiology 18, 1561–1565 (2016).

26 Fontaine, S. S. & Kohl, K. D. The gut microbiota of invasive bullfrog tadpoles responds more rapidly to temperature than a non-invasive congener. Molecular Ecology 29, 2449–2462 (2020).

27 Bestion, E. et al. Climate warming reduces gut microbiota diversity in a vertebrate ectotherm. Nature Ecology & Evolution 1, 0161 (2017).

28 Zhu, L. et al. Environmental temperatures affect the gastrointestinal microbes of the Chinese giant salamander. Frontiers in Microbiology 12, 493 (2021).

29 Moeller, A. H. et al. The lizard gut microbiome changes with temperature and is associated with heat tolerance. Applied and Environmental Microbiology 86, e01181–20 (2020).

30 Kokou, F. et al. Host genetic selection for cold tolerance shapes microbiome composition and modulates its response to temperature. Elife 7, e36398 (2018).

31 Hanage, W. P. Microbiology: microbiome science needs a healthy dose of scepticism. Nature News 512, 247 (2014).

32 Mykles, D. L., Ghalambor, C. K., Stillman, J. H. & Tomanek, L. Grand challenges in comparative physiology: integration across disciplines and across levels of biological organization. Integrative and Comparative Biology 50, 6–16 (2010).

33 Kohl, K. D. A microbial perspective on the grand challenges in comparative animal physiology. Msystems 3, e00146–17 (2018).

34 Pörtner, H.-O., Bock, C. & Mark, F. C. Oxygen-and capacity-limited thermal tolerance: bridging ecology and physiology. Journal of Experimental Biology 220, 2685–2696 (2017).

35 Gangloff, E. J. & Telemeco, R. S. High temperature, oxygen, and performance: Insights from reptiles and amphibians. Integrative and Comparative Biology 58, 9–24 (2018).

36 Gray, K. T., Escobar, A. M., Schaeffer, P. J., Mineo, P. M. & Berner, N. J. Thermal acclimatization in overwintering tadpoles of the green frog, *Lithobates clamitans* (Latreille, 1801). Journal of Experimental Zoology Part A: Ecological Genetics and Physiology 325, 285–293 (2016).

37 Brattstrom, B. H. & Lawrence, P. The rate of thermal acclimation in anuran amphibians. Physiological Zoology 35, 148–156 (1962).

38 Knutie, S. A., Wilkinson, C. L., Kohl, K. D. & Rohr, J. R. Early-life disruption of amphibian microbiota decreases later-life resistance to parasites. Nature Communications 8, 86 (2017).

39 Jani, A. J. & Briggs, C. J. Host and aquatic environment shape the amphibian skin microbiome but effects on downstream resistance to the pathogen *Batrachochytrium dendrobatidis* are variable. Frontiers in Microbiology 9, 487 (2018).

40 Kohl, K. D., Cary, T. L., Karasov, W. H. & Dearing, M. D. Restructuring of the amphibian gut microbiota through metamorphosis. Environmental Microbiology Reports 5, 899–903 (2013).

41 Vences, M. et al. Gut bacterial communities across tadpole ecomorphs in two diverse tropical anuran faunas. The Science of Nature 103, 25 (2016).

42 Fontaine, S. S., Mineo, P. M. & Kohl, K. D. Changes in the gut microbial community of the eastern newt (*Notophthalmus viridescens*) across its three distinct life stages. FEMS Microbiology Ecology 97, fiab021 (2021).

43 Sepulveda, J. & Moeller, A. H. The effects of temperature on animal gut microbiomes. Frontiers in Microbiology 11 (2020).

44 Arango, R. A., Schoville, S. D., Currie, C. R. & Carlos-Shanley, C. Experimental warming reduces survival, cold tolerance, and gut prokaryotic diversity of the eastern subterranean termite, *Reticulitermes flavipes* (Kollar). Frontiers in Microbiology 12, 1116 (2021).

45 Stothart, M. R. et al. Stress and the microbiome: linking glucocorticoids to bacterial community dynamics in wild red squirrels. Biology Letters 12, 20150875 (2016).

46 Zaneveld, J. R., McMinds, R. & Thurber, R. V. Stress and stability: applying the Anna Karenina principle to animal microbiomes. Nature Microbiology 2, 1–8 (2017).

47 Orrock, J. L. & Watling, J. I. Local community size mediates ecological drift and competition in metacommunities. Proceedings of the Royal Society B: Biological Sciences 277, 2185–2191 (2010).

48 Deeg, C. M. et al. *Chromulinavorax destructans*, a pathogen of microzooplankton that provides a window into the enigmatic candidate phylum Dependentiae. PLoS Pathogens 15, e1007801 (2019).

49 Kaboré, O. D., Godreuil, S. & Drancourt, M. Planctomycetes as host-associated bacteria: A perspective that holds promise for their future isolations, by mimicking their native environmental niches in clinical microbiology laboratories. Frontiers in Cellular and Infection Microbiology 10, 729 (2020).

50 Sheremet, A. et al. Ecological and genomic analyses of candidate phylum WPS-2 bacteria in an unvegetated soil. Environmental Microbiology 22, 3143–3157 (2020).

51 Chiang, E. et al. Verrucomicrobia are prevalent in north-temperate freshwater lakes and display class-level preferences between lake habitats. PLoS One 13, e0195112 (2018).

52 Correa, D. T. et al. Multilevel community assembly of the tadpole gut microbiome. Preprint at https://www.biorxiv.org/content/10.1101/2020.07.05.188698v2b (2020).

53 Contijoch, E. J. et al. Gut microbiota density influences host physiology and is shaped by host and microbial factors. Elife 8, e40553 (2019).

54 Trevelline, B. K., Fontaine, S. S., Hartup, B. K. & Kohl, K. D. Conservation biology needs a microbial renaissance: a call for the consideration of host-associated microbiota in wildlife management practices. Proceedings of the Royal Society B 286, 20182448 (2019).

55 Lutterschmidt, W. I. & Hutchison, V. H. The critical thermal maximum: history and critique. Canadian Journal of Zoology 75, 1561–1574 (1997).

56 Gosner, K. L. A simplified table for staging anuran embryos and larvae with notes on identification. Herpetologica 16, 183–190 (1960).

57 Daloso, D. M. The ecological context of bilateral symmetry of organ and organisms. Natural Science 2014 (2014).

58 Goldstein, J. A., Hoff, K. v. S. & Hillyard, S. D. The effect of temperature on development and behaviour of relict leopard frog tadpoles. Conservation Physiology 5, cow075 (2017).

59 Harkey, G. A. & Semlitsch, R. D. Effects of temperature on growth, development, and color polymorphism in the ornate chorus frog *Pseudacris ornata*. Copeia, 1001–1007 (1988).

60 Marian, M. & Pandian, T. Effect of temperature on development, growth and bioenergetics of the bullfrog tadpole *Rana tigrina*. Journal of Thermal Biology 10, 157–161 (1985).

61 Alvarez, D. & Nicieza, A. Effects of temperature and food quality on anuran larval growth and metamorphosis. Functional Ecology 16, 640–648 (2002).

62 Kohl, K. D., Brun, A., Bordenstein, S. R., Caviedes-Vidal, E. & Karasov, W. H. Gut microbes limit growth in house sparrow nestlings (*Passer domesticus*) but not through limitations in digestive capacity. Integrative Zoology 13, 139–151 (2018).

63 Potti, J. et al. Bacteria divert resources from growth for magellanic penguin chicks. Ecology Letters 5, 709–714 (2002).

64 Coates, M. E., Fuller, R., Harrison, G., Lev, M. & Suffolk, S. A comparision of the growth of chicks in the Gustafsson germ-free apparatus and in a conventional environment, with and without dietary supplements of penicillin. British Journal of Nutrition 17, 141–150 (1963).

65 Gaskins, H., Collier, C. & Anderson, D. Antibiotics as growth promotants: mode of action. Animal Biotechnology 13, 29–42 (2002).

66 Hooper, L. V., Littman, D. R. & Macpherson, A. J. Interactions between the microbiota and the immune system. Science 336, 1268–1273 (2012).

67 Gitsels, A., Sanders, N. & Vanrompay, D. Chlamydial infection from outside to inside. Frontiers in Microbiology 10, 2329 (2019).

68 Denver, R. J. Proximate mechanisms of phenotypic plasticity in amphibian metamorphosis. American Zoologist 37, 172–184 (1997).

69 Chevalier, C. et al. Gut microbiota orchestrates energy homeostasis during cold. Cell 163, 1360–1374 (2015).

70 Khakisahneh, S., Zhang, X.-Y., Nouri, Z. & Wang, D.-H. Gut microbiota and host thermoregulation in response to ambient temperature fluctuations. Msystems 5, e00514–20 (2020).

71 Xie, B. et al. *Chlamydomonas reinhardtii* thermal tolerance enhancement mediated by a mutualistic interaction with vitamin B 12-producing bacteria. The ISME Journal 7, 1544–1555 (2013).

72 Gutiérrez-Pesquera, L. M. et al. Testing the climate variability hypothesis in thermal tolerance limits of tropical and temperate tadpoles. Journal of Biogeography 43, 1166–1178 (2016).

73 Litmer, A. R. & Murray, C. M. Critical thermal tolerance of invasion: Comparative niche breadth of two invasive lizards. Journal of Thermal Biology 86, 102432 (2019).

74 Urban, M. C. Accelerating extinction risk from climate change. Science 348, 571–573 (2015).

75 Pearce, T. A. & Paustian, M. E. Are temperate land snails susceptible to climate change through reduced altitudinal ranges? A Pennsylvania example. American Malacological Bulletin 31, 213–224 (2013).

76 Union of Concerned Scientists. Climate change in Pennsylvania: Impacts and solutions for the keystone state (UCS Publications Cambridge, 2008).

77 Huey, R. B. & Kingsolver, J. G. Evolution of thermal sensitivity of ectotherm performance. Trends in Ecology & Evolution 4, 131–135 (1989).

78 Bennett, A. F. Thermal dependence of locomotor capacity. American Journal of Physiology-Regulatory, Integrative and Comparative Physiology 259, R253–R258 (1990).

79 Seebacher, F. & Walter, I. Differences in locomotor performance between individuals: importance of parvalbumin, calcium handling and metabolism. Journal of Experimental Biology 215, 663–670 (2012).

80 Husak, J. F., Fox, S. F., Lovern, M. B. & Bussche, R. A. V. D. Faster lizards sire more offspring: sexual selection on whole-animal performance. Evolution 60, 2122–2130 (2006).

81 Mineo, P. M., Waldrup, C., Berner, N. J. & Schaeffer, P. J. Differential plasticity of membrane fatty acids in northern and southern populations of the eastern newt (*Notophthalmus viridescens*). Journal of Comparative Physiology B 189, 249–260 (2019).

82 Chung, D. J., Sparagna, G. C., Chicco, A. J. & Schulte, P. M. Patterns of mitochondrial membrane remodeling parallel functional adaptations to thermal stress. Journal of Experimental Biology 221, 174458 (2018).

83 Gladwell, R., Bowler, K. & Duncan, C. Heat death in crayfish A*ustropotamobius pallipes*: ion movements and their effects on excitable tissues during heat death. Journal of Thermal Biology 1, 79–94 (1976).

84 Velagapudi, V. R. et al. The gut microbiota modulates host energy and lipid metabolism in mice. Journal of Lipid Research 51, 1101–1112 (2010).

85 Wang, Z. et al. Gut flora metabolism of phosphatidylcholine promotes cardiovascular disease. Nature 472, 57–63 (2011).

86 Pörtner, H. Climate change and temperature-dependent biogeography: oxygen limitation of thermal tolerance in animals. Naturwissenschaften 88, 137–146 (2001).

87 Gräns, A. et al. Aerobic scope fails to explain the detrimental effects on growth resulting from warming and elevated CO_2_ in Atlantic halibut. Journal of Experimental Biology 217, 711–717 (2014).

88 Jutfelt, F. et al. Oxygen-and capacity-limited thermal tolerance: blurring ecology and physiology. Journal of Experimental Biology 221, 169615 (2018).

89 St-Pierre, J., Charest, P.-M. & Guderley, H. Relative contribution of quantitative and qualitative changes in mitochondria to metabolic compensation during seasonal acclimatisation of rainbow trout *Oncorhynchus mykiss*. Journal of Experimental Biology 201, 2961–2970 (1998).

90 Grim, J., Miles, D. & Crockett, E. Temperature acclimation alters oxidative capacities and composition of membrane lipids without influencing activities of enzymatic antioxidants or susceptibility to lipid peroxidation in fish muscle. Journal of Experimental Biology 213, 445–452 (2010).

91 LeMoine, C. M., Genge, C. E. & Moyes, C. D. Role of the PGC-1 family in the metabolic adaptation of goldfish to diet and temperature. Journal of Experimental Biology 211, 1448–1455 (2008).

92 McClelland, G. B., Craig, P. M., Dhekney, K. & Dipardo, S. Temperature-and exercise-induced gene expression and metabolic enzyme changes in skeletal muscle of adult zebrafish (*Danio rerio*). The Journal of Physiology 577, 739–751 (2006).

93 Pichaud, N. et al. Cardiac mitochondrial plasticity and thermal sensitivity in a fish inhabiting an artificially heated ecosystem. Scientific Reports 9, 1–11 (2019).

94 Seebacher, F., Guderley, H., Elsey, R. M. & Trosclair, P. L. Seasonal acclimatisation of muscle metabolic enzymes in a reptile (*Alligator mississippiensis*). Journal of Experimental Biology 206, 1193–1200 (2003).

95 Berner, N. J. & Bessay, E. P. Correlation of seasonal acclimatization in metabolic enzyme activity with preferred body temperature in the Eastern red spotted newt (*Notophthalmus viridescens viridescens*). Comparative Biochemistry and Physiology Part A: Molecular & Integrative Physiology 144, 429–436 (2006).

96 Vigelsø, A., Andersen, N. B. & Dela, F. The relationship between skeletal muscle mitochondrial citrate synthase activity and whole body oxygen uptake adaptations in response to exercise training. International Journal of Physiology, Pathophysiology and Pharmacology 6, 84 (2014).

97 Li, Y., Park, J.-S., Deng, J.-H. & Bai, Y. Cytochrome c oxidase subunit IV is essential for assembly and respiratory function of the enzyme complex. Journal of Bioenergetics and Biomembranes 38, 283–291 (2006).

98 Donohoe, D. R. et al. The microbiome and butyrate regulate energy metabolism and autophagy in the mammalian colon. Cell Metabolism 13, 517–526 (2011).

99 Pryor, G. S. & Bjorndal, K. A. Symbiotic fermentation, digesta passage, and gastrointestinal morphology in bullfrog tadpoles (*Rana catesbeiana*). Physiological and Biochemical Zoology 78, 201–215 (2005).

100 Clark, A. & Mach, N. The crosstalk between the gut microbiota and mitochondria during exercise. Frontiers in Physiology 8, 319 (2017).

101 Payne, N. L. et al. Temperature dependence of fish performance in the wild: links with species biogeography and physiological thermal tolerance. Functional Ecology 30, 903–912 (2016).

102 Van Dijk, P., Tesch, C., Hardewig, I. & Portner, H. Physiological disturbances at critically high temperatures: a comparison between stenothermal Antarctic and eurythermal temperate eelpouts (Zoarcidae). Journal of Experimental Biology 202, 3611–3621 (1999).

103 Schulte, P. M. The effects of temperature on aerobic metabolism: towards a mechanistic understanding of the responses of ectotherms to a changing environment. Journal of Experimental Biology 218, 1856–1866 (2015).

104 Gillooly, J. F., Brown, J. H., West, G. B., Savage, V. M. & Charnov, E. L. Effects of size and temperature on metabolic rate. Science 293, 2248–2251 (2001).

105 Warne, R. W., Kirschman, L. & Zeglin, L. Manipulation of gut microbiota during critical developmental windows affects host physiological performance and disease susceptibility across ontogeny. Journal of Animal Ecology 88, 845–856 (2019).

106 Hoppeler, H. & Weibel, E. R. Scaling functions to body size: theory and facts. Journal of Experimental Biology. 208, 1573–1574 (2005).

107 Hopkins, W. A., Rowe, C. L. & Congdon, J. D. Elevated trace element concentrations and standard metabolic rate in banded water snakes (*Nerodia fasciata*) exposed to coal combustion wastes. Environmental Toxicology and Chemistry: An International Journal 18, 1258–1263 (1999).

108 Sokolova, I. Bioenergetics in environmental adaptation and stress tolerance of aquatic ectotherms: linking physiology and ecology in a multi-stressor landscape. Journal of Experimental Biology 224, 236802 (2021).

109 Sokolova, I. M. & Lannig, G. Interactive effects of metal pollution and temperature on metabolism in aquatic ectotherms: implications of global climate change. Climate Research 37, 181–201 (2008).

110 Peralta-Maraver, I. & Rezende, E. L. Heat tolerance in ectotherms scales predictably with body size. Nature Climate Change 11, 58–63 (2021).

111 Johnston, A. S. et al. Predicting population responses to environmental change from individual-level mechanisms: towards a standardized mechanistic approach. Proceedings of the Royal Society B 286, 20191916 (2019).

112 Trudeau, V. L. et al. Hormonal induction of spawning in 4 species of frogs by coinjection with a gonadotropin-releasing hormone agonist and a dopamine antagonist. Reproductive Biology and Endocrinology 8, 36 (2010).

113 Team, R. C. R: A Language and Environment for Statistical Computing. (2019).

114 Bates, K. A. et al. Amphibian chytridiomycosis outbreak dynamics are linked with host skin bacterial community structure. Nature Communications 9, 693 (2018).

115 Pinheiro, J., et al. Package ‘nlme’. Linear and nonlinear mixed effects models, version 3 (2017).

116 Hulbert, A., Pamplona, R., Buffenstein, R. & Buttemer, W. Life and death: metabolic rate, membrane composition, and life span of animals. Physiological Reviews 87, 1175–1213 (2007).

117 Oksanen, J. et al. Package ‘vegan’. Community ecology package, version 2, 1–295 (2013).

118 Spinazzi, M., Casarin, A., Pertegato, V., Salviati, L. & Angelini, C. Assessment of mitochondrial respiratory chain enzymatic activities on tissues and cultured cells. Nature Protocols 7, 1235–1246 (2012).

119 Singer, J. D. & Willett, J. B. It’s about time: Using discrete-time survival analysis to study duration and the timing of events. Journal of Educational Statistics 18, 155–195 (1993).

120 Wood, S. & Wood, M. S. Package ‘mgcv’. R package version 1, 29 (2015).

121 Caporaso, J. G. et al. Ultra-high-throughput microbial community analysis on the Illumina HiSeq and MiSeq platforms. The ISME Journal 6, 1621 (2012).

122 Bolyen, E. et al. Reproducible, interactive, scalable and extensible microbiome data science using QIIME 2. Nature Biotechnology 37, 852–857 (2019).

123 Callahan, B. J. et al. DADA2: high-resolution sample inference from Illumina amplicon data. Nature Methods 13, 581 (2016).

124 Price, M. N., Dehal, P. S. & Arkin, A. P. FastTree 2–approximately maximum-likelihood trees for large alignments. PloS One 5, e9490 (2010).

125 Quast, C. et al. The SILVA ribosomal RNA gene database project: improved data processing and web-based tools. Nucleic Acids Research 41, D590–D596 (2012).

126 Glassing, A., Dowd, S. E., Galandiuk, S., Davis, B. & Chiodini, R. J. Inherent bacterial DNA contamination of extraction and sequencing reagents may affect interpretation of microbiota in low bacterial biomass samples. Gut Pathogens 8, 1–12 (2016).

127 Eisenhofer, R. et al. Contamination in low microbial biomass microbiome studies: issues and recommendations. Trends in Microbiology 27, 105–117 (2019).

128 Shannon, C. E. A mathematical theory of communication. Mobile Computing and Communications Review 5, 3–55 (2001).

129 Faith, D. P. Conservation evaluation and phylogenetic diversity. Biological Conservation 61, 1–10 (1992).

130 Pielou, E. C. The measurement of diversity in different types of biological collections. Journal of Theoretical Biology 13, 131–144 (1966).

131 Bray, J. R. & Curtis, J. T. An ordination of the upland forest communities of southern Wisconsin. Ecological Monographs 27, 325–349 (1957).

132 Lozupone, C. & Knight, R. UniFrac: a new phylogenetic method for comparing microbial communities. Applied and Environmental Microbiology 71, 8228–8235 (2005).

133 Mallick, H. et al. Multivariable association discovery in population-scale meta-omics studies.Preprint at https://www.biorxiv.org/content/10.1101/2021.01.20.427420v1 (2021).

