## Supplementary Information for "Experimental depletion of gut microbiota diversity reduces host thermal tolerance and fitness under heat stress in a vertebrate ectotherm"

Samantha S. Fontaine[1*], Patrick M. Mineo[2], and Kevin D. Kohl[1]

1. Department of Biological Sciences, University of Pittsburgh, Pittsburgh, PA USA, 15201

2. Biology Department, Elmhurst University, Elmhurst, IL USA, 60126

### Figures

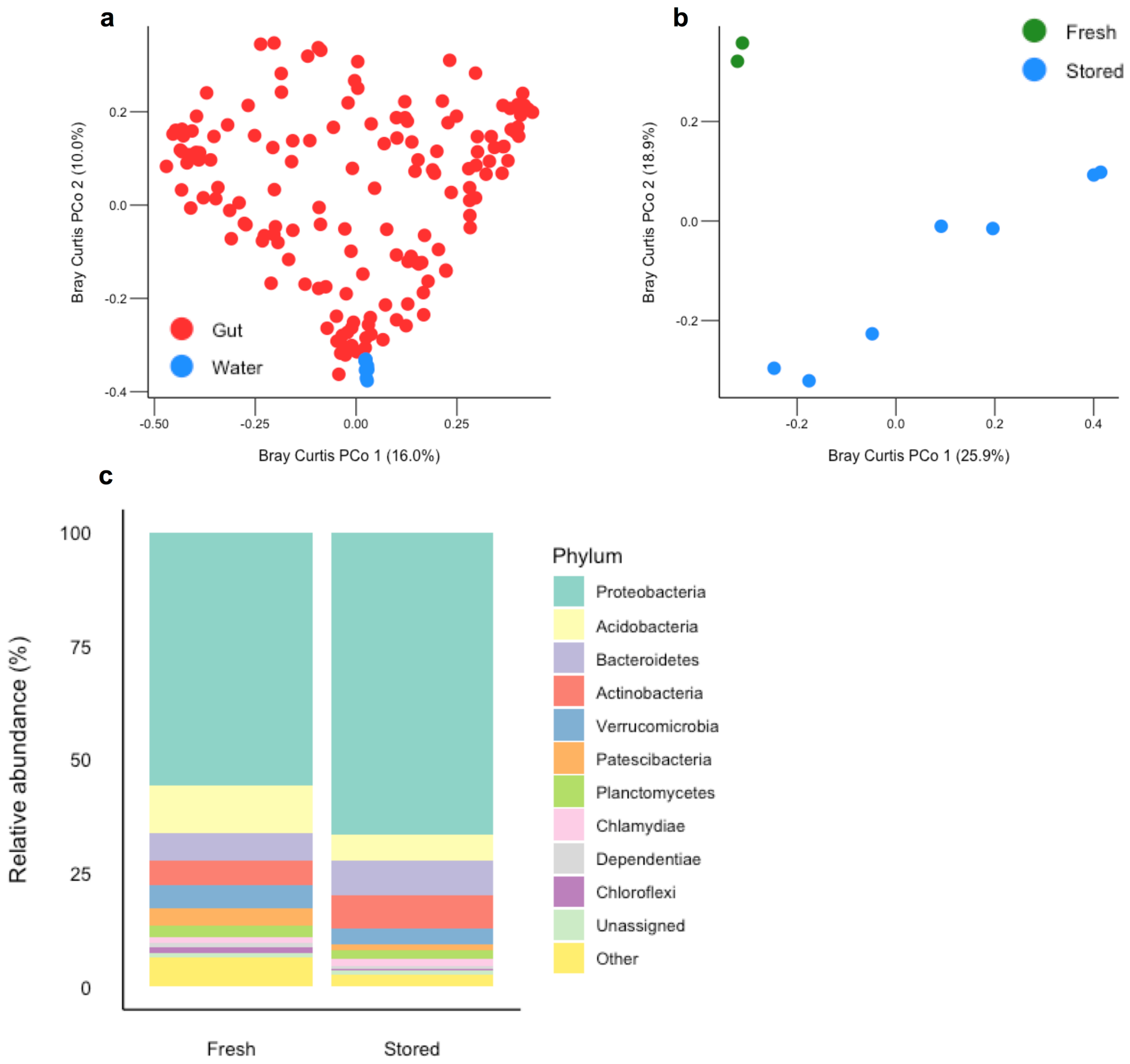


**Supplementary Figure 1. Environmental microbial communities of pond water used for the colonized microbial treatment. a,** Principal Coordinate (PCo) analysis plot based on Bray-Curtis dissimilarity between pond water (both fresh and stored) and tadpole gut microbial community samples. **b,** Principal Coordinate (PCo) analysis plot based on Bray-Curtis dissimilarity between pond water samples collected fresh from the pond or after storage in the laboratory at 4ºC. For both PCoA plots, percentages represent the proportion of variation explained by each axis. **c,** mean relative abundances of bacterial phyla found in pond water samples fresh from the pond or after storage in the laboratory. The top ten most abundant phyla are shown individually, and the remainder are grouped together as “other”. Any bacteria that were unable to be assigned to a phylum are grouped together as “unassigned”.

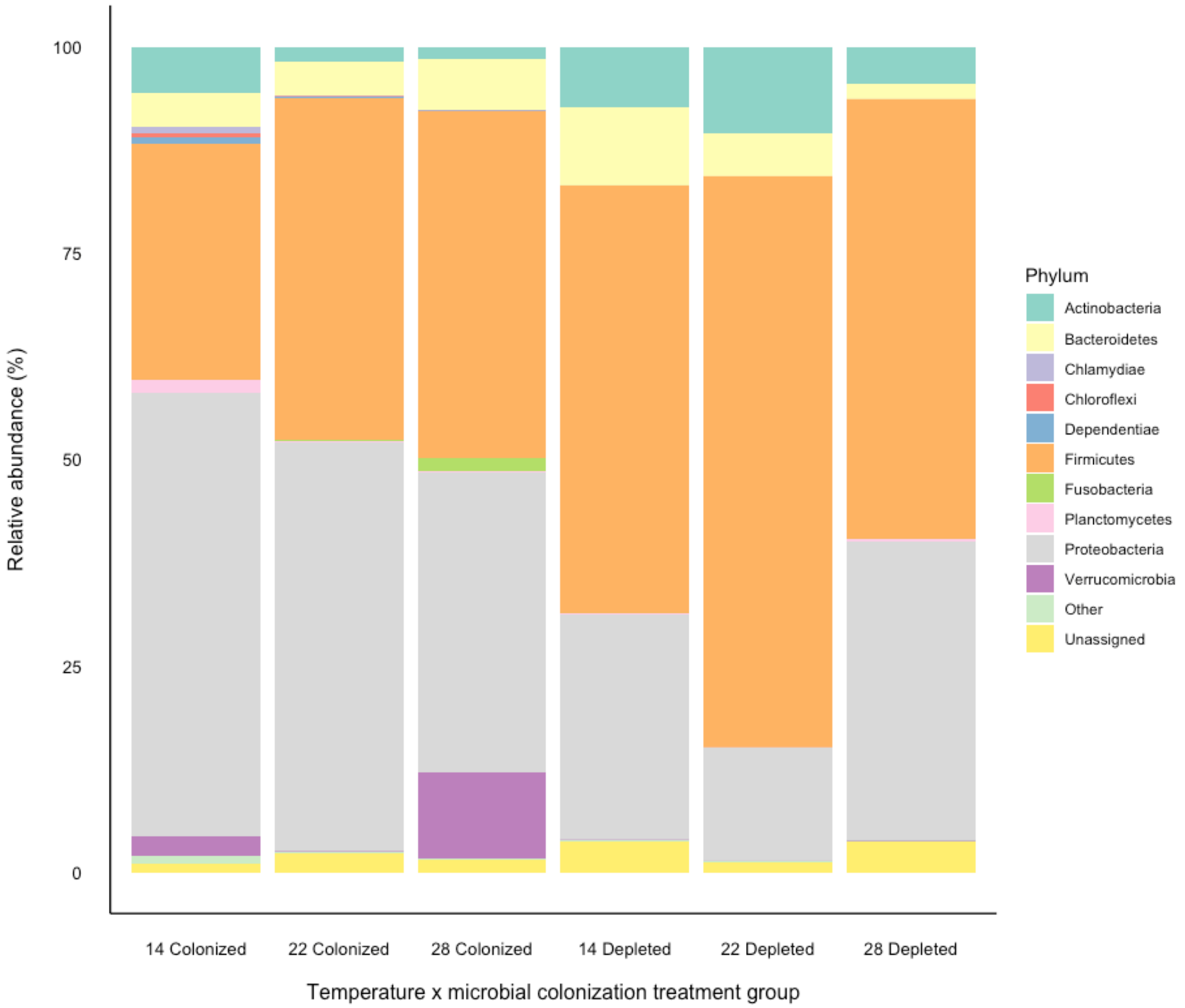


**Supplementary Figure 2. Mean relative abundances of bacterial phyla in gut microbial communities of tadpoles in microbial colonization and acclimation temperature treatment groups.** The top ten most abundant phyla are shown individually, and the remainder are grouped together as “other”. Any bacteria that were unable to be assigned to a phylum are grouped together as “unassigned”.

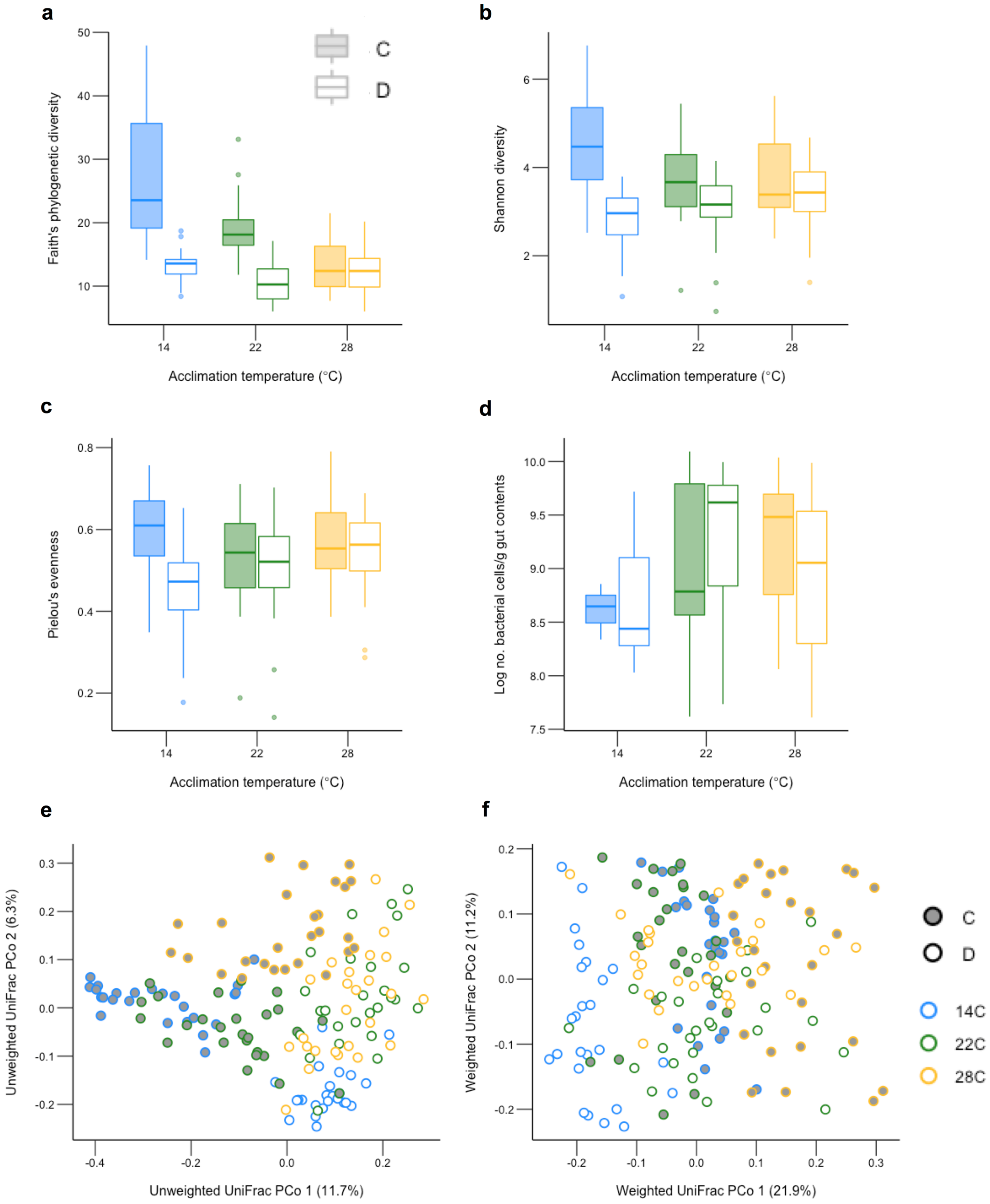


**Supplementary Figure 3. Impacts of microbial colonization treatment and acclimation temperature on tadpole gut microbial communities in experiment 1. a,** Faith's phylogenetic diversity of the gut bacterial community **b,** Shannon diversity of the gut bacterial community **c,** Pielou's evenness within the gut bacterial community **d,** Number of bacterial cells in tadpole gut contents measured using flow cytometry and shown on a log scale **e,** Principal Coordinate (PCo) analysis plot based on Unweighted UniFrac distance between gut bacterial community samples **f,** Principal Coordinate analysis (PCo) plot based on Weighted UniFrac distance between gut bacterial community samples. For boxplots a-c, N = 25-27 animals per group. For boxplot d, N = 9-11 animals per group, except for group 14C colonized where N = 3. For all boxplots, the center line represents the median, the length of the box extends through the IQR, and whiskers extend to 1.5x IQR. All points outside this range are plotted individually. For all principal coordinate analysis plots, percentages represent the proportion of variation explained by each axis. C= colonized tadpoles and D= depleted tadpoles. Colors represent tadpole acclimation temperature.

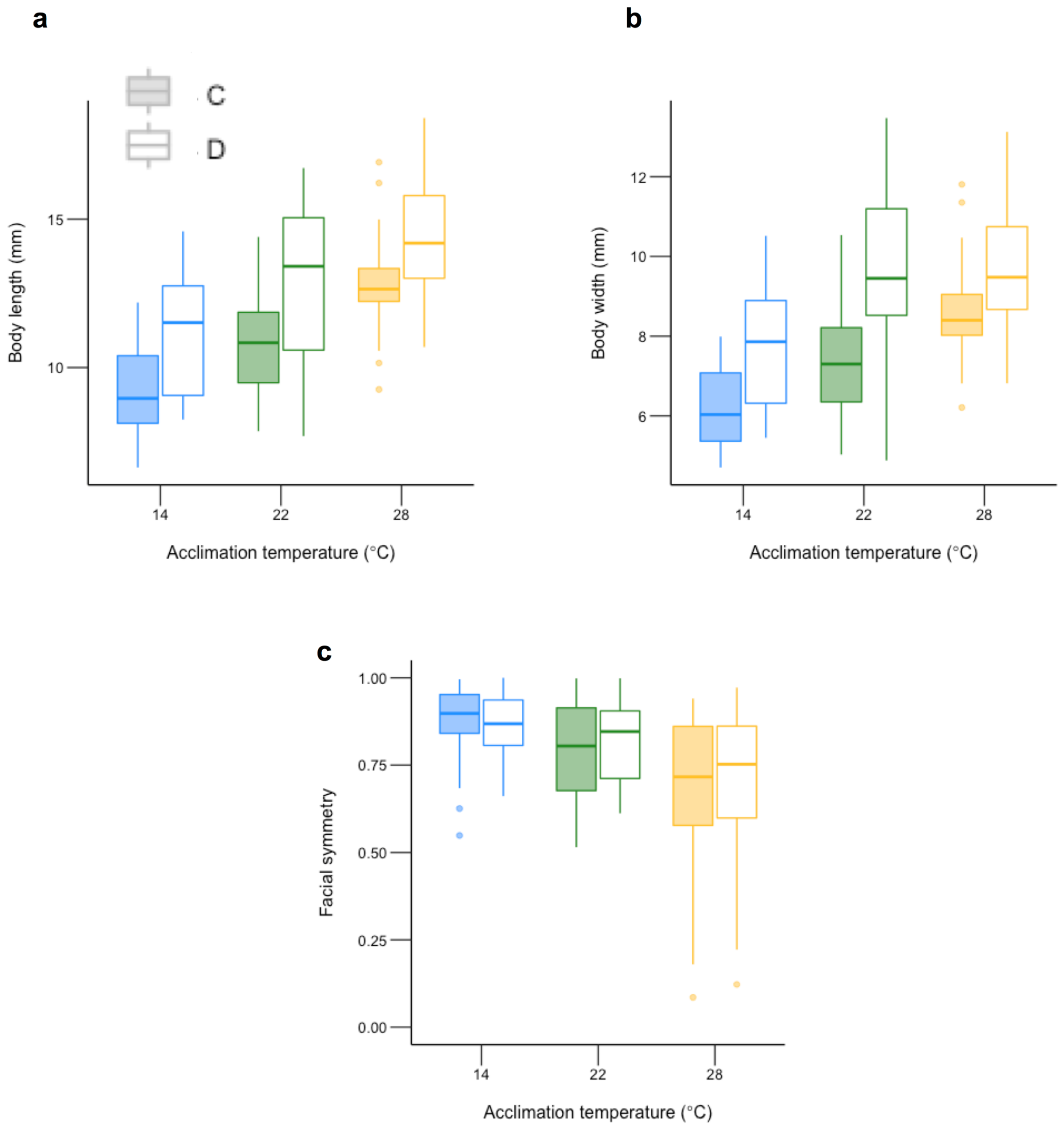


**Supplementary Figure 4. Tadpole morphometrics across microbial colonization and acclimation temperature treatment groups from experiment 1. a,** tadpole body length **b,** tadpole body width **c,** tadpole facial symmetry, calculated as the absolute value, subtracted from 1, of the difference between the distance from the center of each eye to the tip of the nose. The center line of each boxplot represents the median, the length of the box extends through the IQR, and whiskers extend to 1.5x IQR. All points outside this range are plotted individually. C= colonized tadpoles and D= depleted tadpoles. Colors represent tadpole acclimation temperature. N = 27 animals per group.

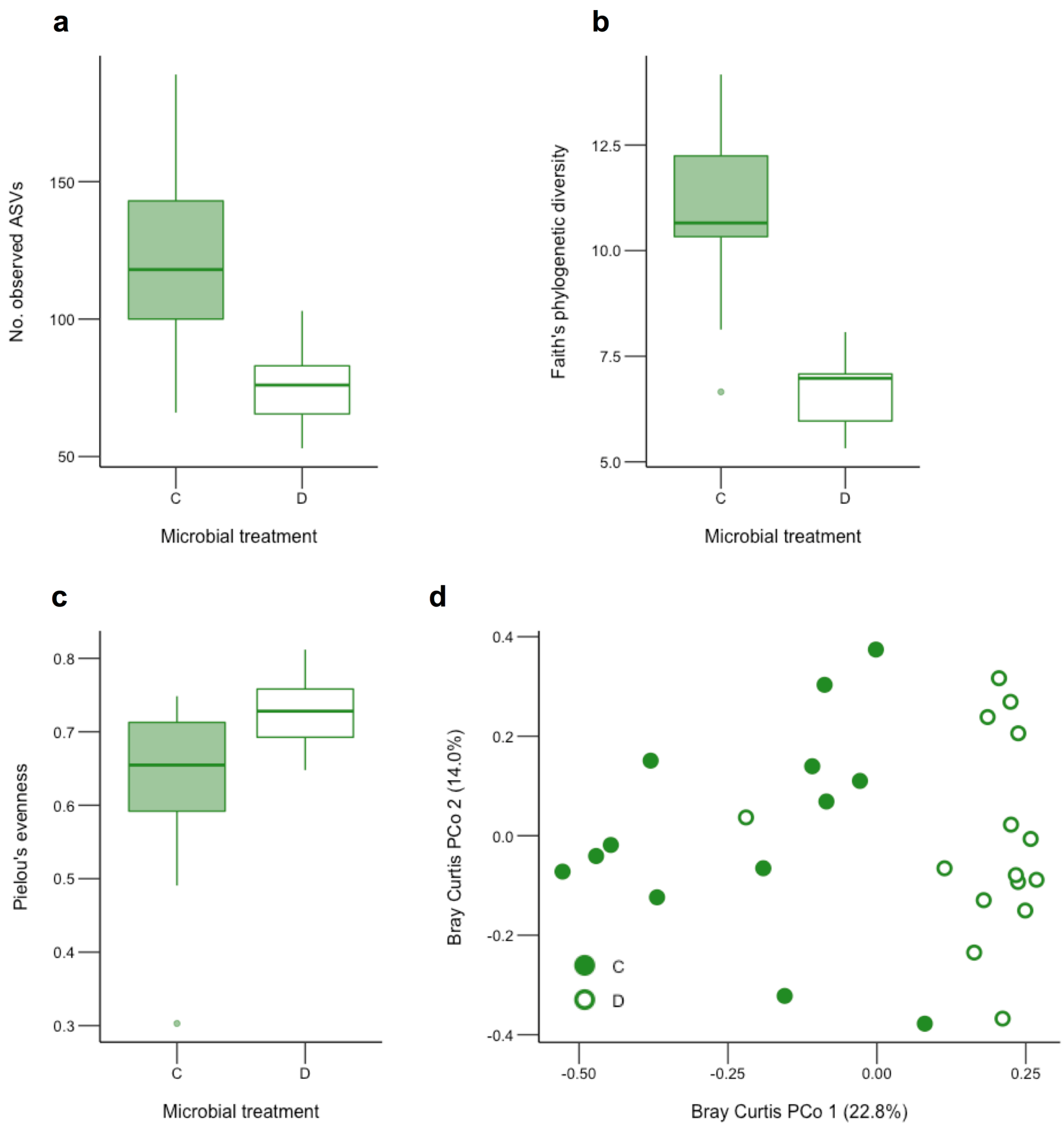


**Supplementary figure 5. Impact of microbial colonization treatment on tadpole gut microbial communities in experiment 2. a,** Number of bacterial ASVs in the tadpole gut bacterial community, **b,** Faith's phylogenetic diversity of the tadpole gut bacterial community, **c,** Pielou's evenness of the tadpole gut bacterial community, and **d,** Principal Coordinate (PCo) analysis plot based on Bray-Curtis dissimilarity between gut bacterial community samples. Percentages represent the proportion of variation explained by each axis. The center line of each boxplot represents the median, the length of the box extends through the IQR, and whiskers extend to 1.5x IQR. All points outside this range are plotted individually. C= colonized tadpoles and D= depleted tadpoles. N = 17 animals per group.

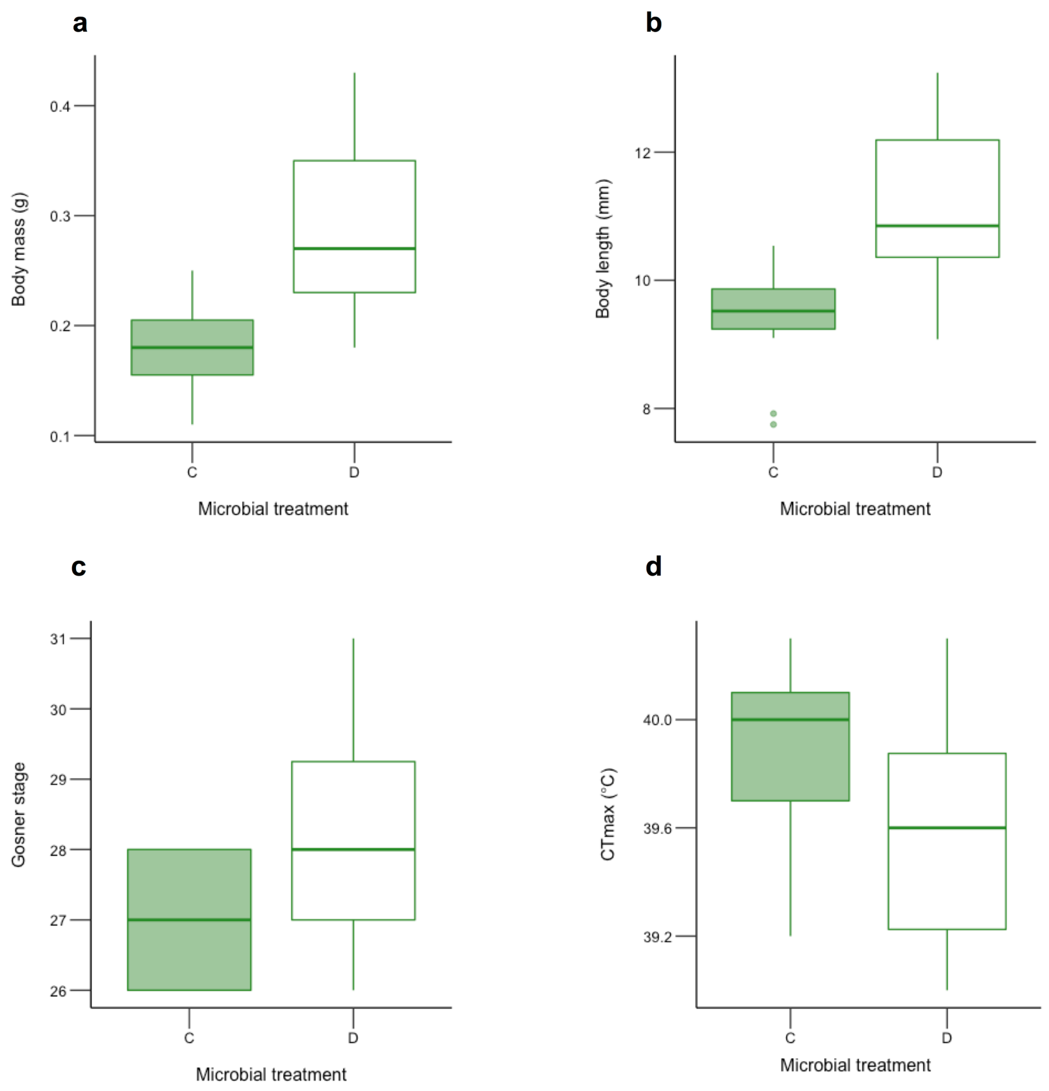


**Supplementary Figure 6. The impact of microbial colonization treatment on tadpole morphometrics and heat tolerance in experiment 2. a,** tadpole body mass **b,** tadpole body length **c,** tadpole developmental stage based on the Gosner system **d,** tadpole critical thermal maxima ($CT_{max}$). The center line of each boxplot represents the median, the length of the box extends through the IQR, and whiskers extend to 1.5x IQR. All points outside this range are plotted individually. C= colonized tadpoles and D= depleted tadpoles. N = 17-20 animals per group.

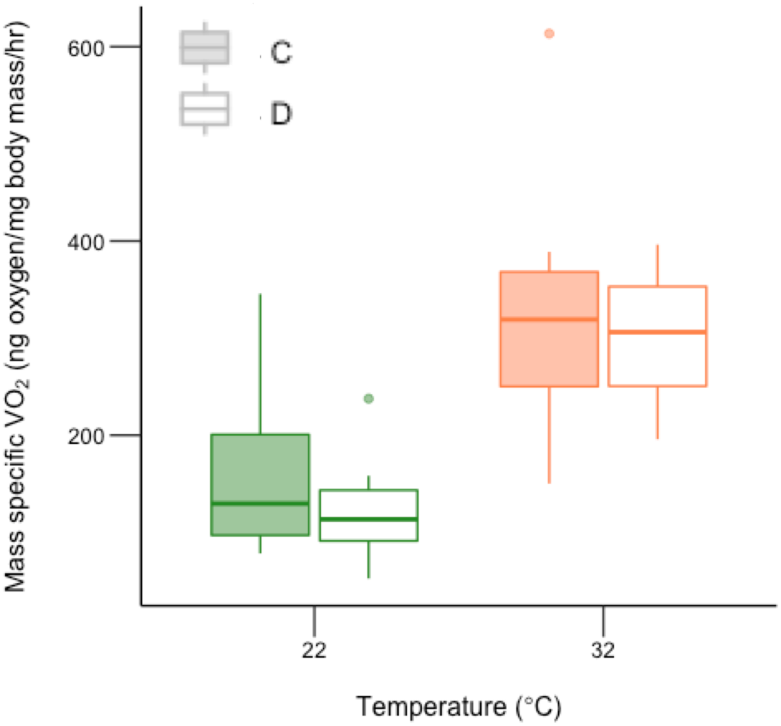


**Supplementary Figure 7. Impacts of microbial colonization treatment and assay temperature on tadpole mass-specific resting metabolic rate.** The center line of each boxplot represents the median, the length of the box extends through the IQR, and whiskers extend to 1.5x IQR. All points outside this range are plotted individually. C= colonized tadpoles and D= depleted tadpoles. On the y-axis $VO_2$ refers to oxygen consumption. N = 9-10 animals per group.

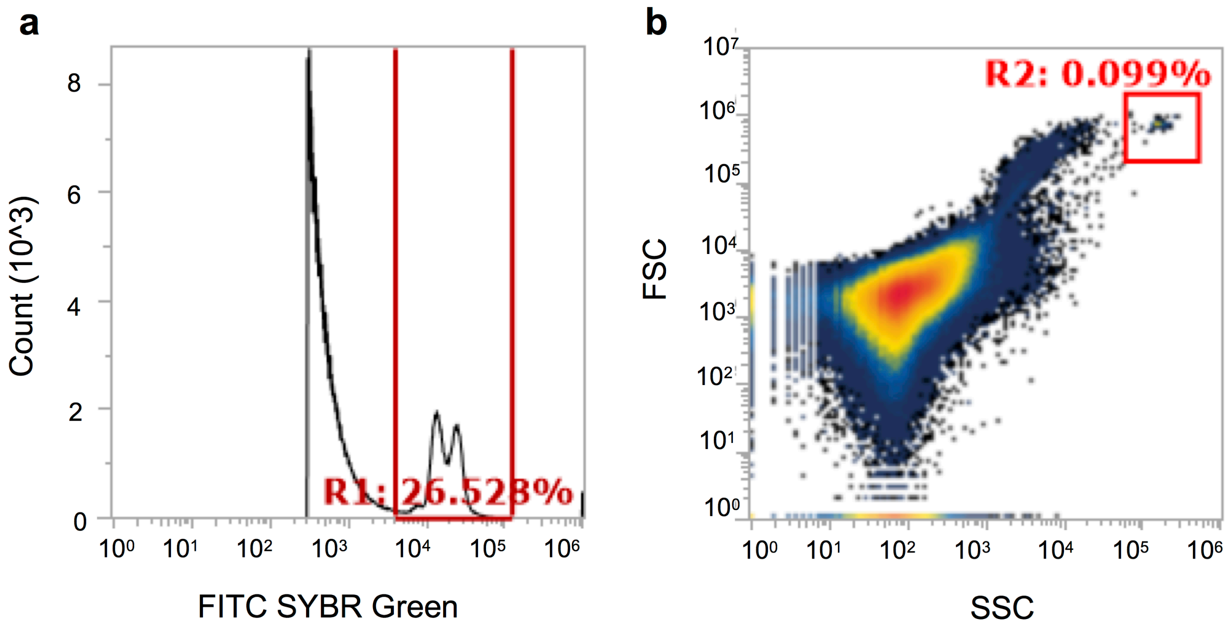


**Supplementary Figure 8. Plots used to determine gating parameters for flow cytometry from one representative sample. a,** A plot of fluorescein isothiocyanate (FITC) vs. cell counts was used to distinguish cells stained with SYBR Green dye from all other cells **b,** A plot of forward scatter (FSC) vs. side scatter (SSC) was used to distinguish populations of counting beads from all other events. To establish initial gates for counting beads, blank samples spiked only with beads were used. On both plots, red rectangles represent the gates and surround the events counted. Percentages indicate the proportion of events within the gate out of total events. R1= stained bacterial cells and R2=counting beads.

### Tables

**Supplementary Table 1. Results of statistical tests examining the effects of acclimation temperature, microbial colonization treatment, and their interaction on tadpole gut microbial community alpha and beta diversity.** NS= non-significant. N/A= non-applicable, an interactive effect could not be measured for this test. For experiment 1 models, tadpole tank was included as a random effect. A single acclimation temperature and individual housing was used in experiment 2 so no effects of temperature were tested, and tank was not included in models. Significant effects are shown in bold.

| **Diversity metric** | **Temperature effect (P, test statistic)** | **Colonization effect (P, test statistic)** | **T x C interactive effect (P, test statistic)** |
|---|---|---|---|
| *Experiment 1* | | | |
| *Alpha diversity (GLMMs)* | | | |
| No. observed ASVs | **<0.001, 33.8** | **<0.001, 67.0** | **<0.001, 38.2** |
| Faith's phylogenetic diversity | **<0.001, 54.0** | **<0.001, 71.4** | **<0.001, 40.5** |
| Shannon diversity | NS | **<0.001, 34.3** | **<0.001, 16.4** |
| Pielou's evenness | NS | **<0.001, 12.0** | **0.02, 5.9** |
| *Beta diversity (PERMANOVAs)* | | | |
| Bray-Curtis | **<0.001, 9.6** | **<0.001, 19.2** | **<0.001, 4.1** |
| Unweighted UniFrac | **<0.001, 5.1** | **<0.001, 12.8** | **<0.001, 3.9** |
| Weighted UniFrac | **<0.001, 8.3** | **<0.001, 14.4** | **<0.001, 5.9** |
| *Beta dispersion (PERMDISPs)* | | | |
| Bray-Curtis | **<0.01, 6.1** | **<0.001, 20.8** | N/A |
| Unweighted UniFrac | **<0.001, 19.2** | **<0.001, 37.4** | N/A |
| Weighted UniFrac | **<0.01, 7.1** | **<0.01, 8.0** | N/A |
| *Experiment 2* | | | |
| *Alpha diversity (Kruskal- Wallis)* | | | |
| No. observed ASVs | N/A | **<0.001, 15.0** | N/A |
| Faith's phylogenetic diversity | N/A | **<0.001, 16.2** | N/A |
| Shannon diversity | N/A | NS | N/A |
| Pielou's evenness | N/A | **<0.01, 7.8** | N/A |
| *Beta diversity (PERMANOVAs)* | | | |
| Bray-Curtis | N/A | **<0.001, 5.2** | N/A |
| Unweighted UniFrac | N/A | **<0.001, 8.0** | N/A |
| Weighted UniFrac | N/A | **<0.001, 4.5** | N/A |
| *Beta dispersion (PERMDISPs)* | | | |
| | N/A | NS | N/A |
| Bray-Curtis | N/A | **<0.05, 5.7** | N/A |
| Unweighted UniFrac | N/A | NS | N/A |
| Weighted UniFrac | | | |

**Supplementary Table 2. Relative abundances of bacterial phyla and genera in tadpole gut microbial communities that were significantly impacted by microbial colonization treatment or acclimation temperature.** Relative abundances are displayed as means ± s.e.m. The group in which the specific taxa was most abundant is displayed in bold. Statistical testing was conducted using MaAsLin2. N.O=not observed. P-values were corrected using the FDR method.

| | Relative abundance (%) | | | FDR P-value | Coefficient |
|---|---|---|---|---|---|
| **Temperature** | **14°C** | **22°C** | **28°C** | | |
| **Phyla** | | | | | |
| Acidobacteria | **0.10 ± 0.03** | 0.01 ± <0.01 | <0.01 | 0.03 | -0.04 |
| **Colonization** | **Colonized** | **Depleted** | | | |
| **Phyla** | | | | | |
| Chlamydiae | **0.33 ± 0.07** | <0.01 | | <0.01 | -0.04 |
| Chloroflexi | **0.20 ± 0.05** | 0.02 ± 0.01 | | 0.03 | -0.02 |
| Dependentiae | **0.32 ± 0.07** | <0.01 | | <0.01 | -0.04 |
| Plantomycetes | **0.62 ± 0.13** | 0.11 ± 0.03 | | 0.01 | -0.04 |
| Proteobacteria | **46.76 ± 2.77** | 26.13 ± 2.68 | | <0.01 | -0.27 |
| WPS-2 | **0.05 ± 0.01** | N.O. | | 0.02 | -0.01 |
| Firmicutes | 37.83 ± 2.69 | **59.13 ± 2.63** | | <0.01 | 0.25 |
| **Genera** | | | | | |
| *Aquicella* | **0.52 ± 0.13** | N.O. | | <0.01 | -0.05 |
| *Aquisphaera* | **0.17 ± 0.04** | <0.01 | | 0.01 | -0.03 |
| *Coxiella* | **0.08 ± 0.02** | <0.01 | | 0.01 | -0.02 |
| *Roseiarcus* | **0.56 ± 0.13** | N.O. | | <0.01 | -0.05 |
| *Singulisphaera* | **0.13 ± 0.33** | N.O. | | <0.01 | -0.02 |
| Uncultured *Diplorickettsiaceae* | **0.21 ± 0.05** | <0.01 | | 0.01 | -0.03 |
| Uncultured *Simkaniaceae* | **0.12 ± 0.02** | <0.01 | | 0.01 | -0.02 |
| Uncultured *Vermiphilaceae* | **0.09 ± 0.03** | <0.01 | | 0.01 | -0.02 |
| *Xanthobacter* | **8.49 ± 1.21** | 1.46 ± 0.45 | | <0.01 | -0.17 |
| *Bacillus* | 6.43 ± 1.35 | **31.81 ± 2.85** | | <0.01 | 0.43 |
| *Rikenella* | 1.59 ± 0.35 | **5.45 ± 1.02** | | 0.03 | 0.13 |

**Supplementary Table 3. Results of statistical tests examining the effects of acclimation temperature, microbial colonization treatment, and their interaction on tadpole morphometrics in experiments 1 and 2.** For experiment 1 models, tadpole tank was included as a random effect in all models, and body length was included as a covariate in the facial symmetry model. NS= non-significant. N/A= non-applicable. A single acclimation temperature and individual housing was used in experiment 2 so no effects of temperature were tested, and tank was not included in models. Significant effects are shown in bold.

| Metric | Temperature effect (P, test statistic) | Colonization effect (P, test statistic) | T x C interactive effect (P, test statistic) |
|---|---|---|---|
| *Experiment 1 (GLMMs)* | | | |
| Body mass (g) | **<0.001, 46.2** | **<0.001, 43.1** | NS |
| Body length (mm) | **<0.001, 72.4** | **<0.001, 27.6** | NS |
| Body width (mm) | **<0.001, 53.9** | **<0.001, 40.7** | NS |
| Gosner stage | **<0.001, 85.7** | **<0.001, 36.6** | **<0.01, 12.1** |
| Facial symmetry | **<0.001, 34.3** | NS | NS |
| *Experiment 2 (GLMs)* | | | |
| Body mass (g) | N/A | **<0.001, 26.3** | N/A |
| Body length (mm) | N/A | **<0.001, 27.7** | N/A |
| Gosner Stage | N/A | **<0.001, 13.1** | N/A |

**Supplementary Table 4. Relative abundances of phospholipid species that differed between colonized and depleted tadpoles.** Relative abundances are displayed as means ± s.e.m. The group in which a species was most abundant is shown in bold. Significance was determined using ANOVAs and the response screening function in JMP. P-values were corrected using the FDR method. Phospholipid species nomenclature is depicted as “phospholipid class (no. carbons: no. double bonds)”. PC= phosphatidylcholine, PE= phosphatidylethanolamine, PS= phosphatidylserine, PI= phosphatidylinositol.

| | Relative abundance (%) | | FDR P-value | F statistic |
|---|---|---|---|---|
| | Colonized | Depleted | | |
| **Phospholipid Species** | | | | |
| PC(35:2) | **0.71 ± 0.06** | 0.48 ± 0.04 | 0.01 | 11.41 |
| PC(36:1) | **1.01 ± 0.06** | 0.68 ± 0.05 | <0.01 | 18.03 |
| PC(37:2) | **0.15 ± 0.02** | 0.09 ± 0.01 | 0.02 | 10.48 |
| PC(37:6) | **0.12 ± 0.01** | 0.06 ± 0.01 | <0.01 | 19.32 |
| PC(38:6) | **0.55 ± 0.08** | 0.43 ± 0.03 | 0.01 | 11.11 |
| PE(29:1) | **1.50 ± 0.11** | 1.00 ± 0.15 | 0.04 | 8.26 |
| PS(42:9) | **0.12 ± 0.02** | 0.05 ± 0.01 | 0.01 | 11.98 |
| PS(44:12) | **0.11 ± 0.02** | 0.05 ± 0.01 | 0.03 | 8.97 |
| PC(34:3) | 1.54 ± 0.06 | **2.12 ± 0.15** | 0.02 | 10.24 |
| PE(32:0) | 0.22 ± 0.01 | **0.31 ± 0.02** | 0.01 | 10.89 |
| PI(36:4) | 0.08 ± 0.01 | **0.12 ± 0.01** | <0.01 | 12.90 |
| PI(38:5) | 0.37 ± 0.04 | **0.61 ± 0.05** | <0.01 | 15.51 |
